# Phage Endolysin Enables Targeted Manipulation of the Small Intestinal Microbiota and Uncovers Niche Overlap Between Oral and Butyrate-Producing Taxa

**DOI:** 10.64898/2026.01.06.697609

**Authors:** Julian R. Garneau, Aline Altenried, Youzheng Teo, Xiaobing Wu, Sarah McHugh, Michael Taschner, Carla Hernández-Cabanyero, Johann Mignolet, Pénélope Gomes, Hugues De Villiers de la Noue, Grégory Resch, Pascale Vonaesch

## Abstract

Oral bacterial overgrowth in the small intestine has been associated with dysbiosis, impaired nutrient absorption, and stunted growth in undernourished children, a condition referred to as small intestinal oral bacterial overgrowth (SIOBO). Here, we explore the use of a phage-derived lysin as a precision antimicrobial to selectively target *Streptococcus salivarius* within complex microbial communities. Through a newly developed, medium-throughput bioinformatic and wet-lab pipeline we identified, cloned and produced a prophage-encoded lysin from *S. salivarius* and demonstrated its potent and specific lytic activity against a panel of 49 clinical *S. salivarius* strains from stunted children, while sparing related species such as *S. mitis*, *S. parasanguinis and S. thermophilus*. Application of the lysin to human stool-derived *in vitro* communities and to mice colonized with *S. salivarius* led to an approximate 2-3 log reduction in *S. salivarius* abundance while preserving overall bacterial community composition. However, we observed a negative correlation between *S. salivarius* and *Coprococcus comes in vitro*, and *Eubacterium xylanophilum, Akkermansia, Lactobacillus and Ruminococcus in vivo*. Spent medium assays confirmed niche overlap between 10 clinical strains of *S. salivarius* and 28 different taxa involved in butyrate production. Together, these results suggest that phage-derived lysins can offer multiple benefits by selectively removing ectopically colonized oral taxa and indirectly promoting beneficial anaerobes.

## INTRODUCTION

The small intestinal microbiota is increasingly recognized as a critical factor in the pathophysiology of non-communicable diseases^1–7^. In several such conditions, the disproportionate expansion of specific oral bacterial taxa has been associated with gut microbial dysbiosis and host dysfunction^2^. One striking example is childhood stunting, which is defined as a height-for-age two standard deviations below the WHO standard curve and affects approximately 149 million children worldwide^3,8,9^. Associated with impaired physical growth and cognitive development with long-lasting effects, stunting remains largely unresponsive to nutritional interventions alone^8,10,11^. Recent findings have implicated a reduction in butyrate producers^3,12^ as well as small intestinal oral bacterial overgrowth (SIOBO) as a contributor to the chronic intestinal inflammation frequently observed in stunted children^3,7,13,14^. Notably, ectopic colonization of the small intestine by oral bacteria was reported in nearly 80% of affected individuals, with *Streptococcus salivarius* emerging as the most dominant species^13^.

Although *S. salivarius* is often regarded as a commensal, its overabundance in the small intestines of stunted children suggests a potentially detrimental role in this context. Previous studies have linked elevated prevalence and abundance of *S. salivarius* to the exacerbation of inflammatory conditions such as rhinitis^15^. Also, a negative correlation has been observed between its absolute abundance in the duodenum and normal child growth^7^. High levels of this species on tissue culture cells or in the murine small intestine has been shown to impair lipid absorption into enterocytes^3^. Together, these findings support the rationale for microbiota-targeted strategies in preventing stunting, specifically those aimed at reducing the burden of potentially harmful oral bacteria like *S. salivarius*.

Targeted approaches that spare the overall microbiota could alleviate intestinal inflammation and its downstream consequences, including compromised nutrient uptake. Such precision interventions may contribute to restoring a healthier microbial ecosystem. Given their capacity to target specific species, we first hypothesised that bacteriophages could be leveraged to selectively reduce the overabundance of *S. salivarius* in the small intestine of stunted children, while preserving the overall structure and function of the resident microbiota. Indeed, phages are highly species-specific bacterial predators and can enable the selective targeting of opportunistic pathogens, offering a means to modulate host physiology with greater precision^16,17^. This was highlighted in a study on alcohol-related liver disease, where a cocktail of virulent phages targeting cytolysin-producing *Enterococcus faecalis* successfully reversed disease in mice by reducing bacterial abundance^18^. Importantly, these findings also showed that partial reduction of a disease-associated commensal, rather than its complete elimination, may suffice to confer therapeutic benefit.

To date, only a single *S. salivarius* prophage, YMC-2011 has been described in the literature. The prophage was induced using mitomycin C and visualized by transmission electron microscopy. However, the authors were unable to obtain infection plaques, propagate the phage, or purify it^19^. Thus, no lytic phages have yet been characterized for this species, limiting therapeutic development using monophage preparations or phage cocktails. In addition, the use of phages in complex host environments requires careful evaluation of factors such as their stability, host accessibility, potential bacterial resistance development, innate bacterial defence systems, and the risk of undesired horizontal gene transfer^20–22^. While these barriers can be overcome, they underscore the importance of pursuing complementary or alternative strategies, such as the use of phage lysins, which offers the possibility of selectively modulating microbial communities in the context of non-communicable diseases^23^.

Phage lysins are considered less prone to resistance development, making them attractive candidates for broad-scale therapeutic applications^24–26^. Also, owing to their recent notable clinical development, they are increasingly recognized as a leading alternative to traditional antibacterial therapies^27^. While no lysin has yet received clinical approval^28^, several candidates are advancing through preclinical and clinical development, particularly targeting methicillin-resistant *Staphylococcus aureus* (MRSA)^29,30^, *Pseudomonas aeruginosa*^31,32^ and, particularly relevant for the present study, various *Streptococcal* species such as *Streptococcus pneumoniae*^33,34^ and *Streptococcus agalactiae*^35^. Recent phase 2 and 3 randomized clinical trials with the *S. aureus* phage lysin highlighted its favourable safety profile following intravenous administration^36,37^.

In this study, following full bacterial genome sequencing and annotation, we identified two novel prophages in the genomes of *S. salivarius* AF151 and AF238. We subsequently cloned and functionally characterized one of their encoded lysin, termed Lys_AF151 and show its specific lytic activity against a representative panel of clinical *S. salivarius* isolates in monoculture, within complex stool-derived *in vitro* communities as well as a *in vivo* murine model of *S. salivarius* intestinal colonization. Most interestingly, by reducing their niche competitor *S. salivarius*, Lys_AF151 also increased the level of butyrate-producing strains, thus overall leading to a normalization of the intestinal ecosystem within the stunted samples/individuals.

Overall, this study highlights the potential of lysins as precision antimicrobials capable of selectively reducing target taxa. We also demonstrate their value as functional tools for exploring microbial ecology, since shifts in non-target species after target depletion can indirectly reveal underlying interactions and dependencies.

## RESULTS

### *Streptococcus salivarius* clinical strains AF151 and AF238 harbour functional prophages encoding for endolysins of the glycoside hydrolase family

To identify *S. salivarius* phage encoded endolysins, we first searched for chromosome-integrated or episomal prophage sequences in our collection of clinical *S. salivarius* isolates. Using PHASTEST phage prediction tool^38^, we identified two prophage regions in the AF151 and AF238 strains annotated as complete (Phi_Aura_AF151 and Phi_Alten_AF238, respectively). We first sought to determine whether mitomycin C stress could induce prophage activation and virion production. Transmission electron microscopy (TEM) performed on the 0.22 μm filtered *S. salivarius* mitomycin C-induced supernatants confirmed comparative genomics results classifying Phi_Aura_AF151 and Phi_Alten_AF238 as *Siphoviruses* (Supp. Fig. 1). Furthermore, sequencing of the filtered supernatant revealed pronounced enrichment of sequencing reads mapping to the prophage regions on the bacterial genomes (Fig 1A). Taken together, these results confirm that the two identified prophage regions are functional and can lead to the production of *Siphoviruses* equipped with a headful DNA packaging system^39^, as revealed by PhageTerm^40^ analysis (Fig. 1B). The two phage genomes were of 43’633 bp and 43’073 bp in length, respectively. Analysis of genes annotations revealed a gene encoding for a glycoside hydrolase in each of the two prophages (Fig. 1C). We further analyzed the predicted endolysin sequences using InterProScan^41^ and identified a glycosyl hydrolase domain along with cell wall and choline-binding domains (Fig. 1D). Since both endolysin sequences were highly similar, with 86% nucleotide identity over 100% coverage, and 96% identity at the amino acid level, we selected Phi_Aura_AF151 lysin (Lys_AF151) for cloning and overexpression in *E. coli*. Lys_AF151 was successfully cloned, overexpressed and purified yielding a final concentration of 10 mg/mL (Fig. 1E).

**Figure 1.**
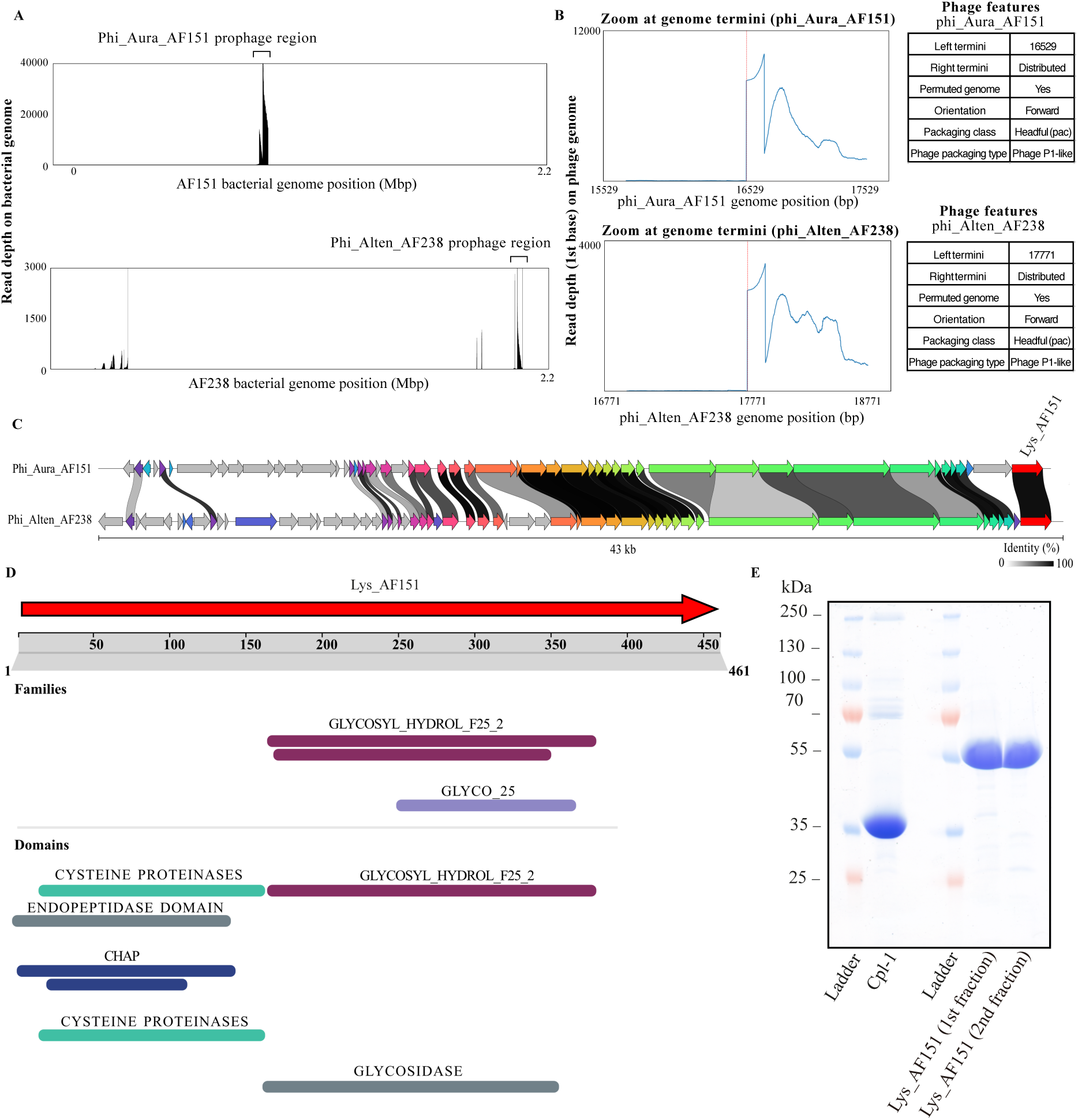
Induction and genomic comparison of prophages Phi_Aura_AF151 and Phi_Alten_AF238 following mitomycin C treatment. (A) Read coverage along the bacterial chromosome after mitomycin C-induced prophage excision. The graph shows the coverage depth of viral reads mapped to the AF151 and AF238 genomes, with a clear enrichment peak at the prophage region, indicating successful induction of Phi_Aura_AF151 and Phi_Alten_AF238. (B) Genome termini analysis and gDNA packaging strategy of induced phages with plots show read-start depth near the predicted genome termini of two phages. The accumulation of sequencing reads-start at the left termini (red vertical line) indicates signatures of physical ends of the packaged phage genomes inside phage capsid. Both phages exhibit a typical signature of headful packaging (pac-type), with a sharp left terminus and distributed right ends, consistent with a permuted genome structure. (C) Genomic alignment of the two induced and assembled phage genomes. Arrows represent annotated open reading frames (ORFs), given an identical color if protein had high identity between two genomes. Grey shading between genomes indicates regions of nucleotide identity, with darker shades corresponding to higher protein identity. Lysin genes are highlighted in red to indicate their position and conservation across the two phages. (D) InterProScan domain annotation and of the Lys_AF151 lysin. The domain map shows predicted functional motifs across the protein sequence, indicating conserved catalytic and cell wall binding regions. E) SDS-PAGE gel showing the expression and purification profiles of the Lys_AF151 and Cpl-1 lysins. 10 uL of protein samples were loaded on gel and visualized on a 12% acrylamide gel.

### Prophage endolysin Lys_AF151 exhibits potent and specific lytic activity against a variety of clinical *S. salivarius* strains

To evaluate the activity and host specificity of the Lys_AF151 lysin, we tested 49 clinical isolates of *S. salivarius*, 10 of *S. parasanguinis*, nine of *S. mitis*, and 2 of *S. thermophilus* (Supp. Table 1) using both an adapted disk diffusion assay (Fig. 2A) and a liquid broth inhibition assay (Fig. 2B, Supp. Fig. 2 and Supp. Table 1). Lys_AF151 showed clear inhibitory activity against 46 out of 49 *S. salivarius* isolates (94%), with varying degrees of inhibition depending on the strain. No inhibition was observed against any *S. mitis* or *S. thermophilus* isolates, and only three of 10 S. *parasanguinis* isolates displayed slight susceptibility, supporting the high specificity of Lys_AF151 for *S. salivarius*. As a negative control, we tested Cpl-1, a phage lysin known to target *S. pneumoniae*^34^, produced and purified using the same procedure as Lys_AF151, which had no effect on the growth of *S. salivarius* strains. For strains tested in agar and liquid, the results from both assays were overall highly concordant with minor differences in the inhibition strength by assay type for a few of the strains tested (Fig. 2C). For 2 strains we observed lysin activity on agar but detected no effect in the broth assay.

**Figure 2.**
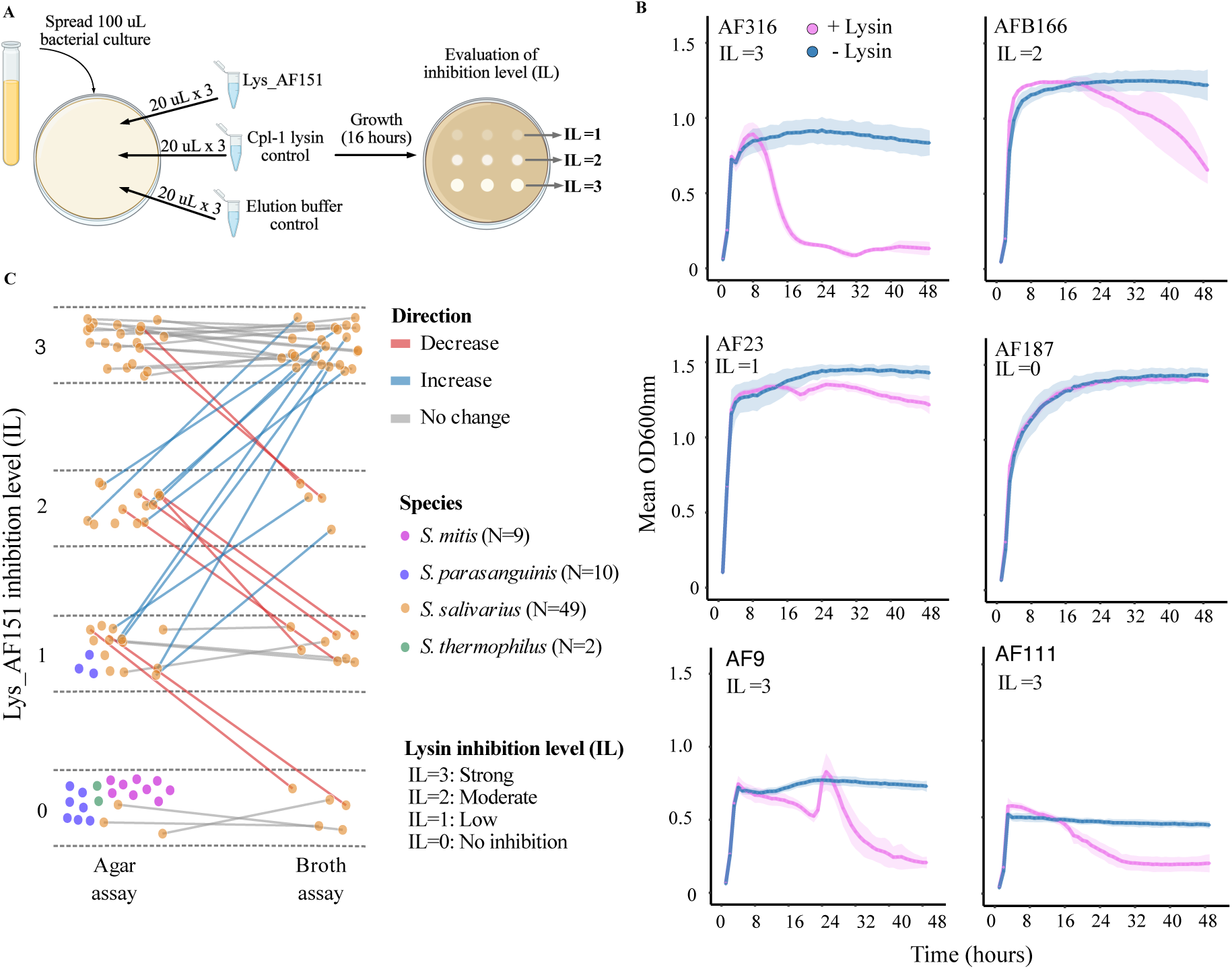
Inhibition of *Streptococcus* isolates by endolysin Lys_AF151 in agar and broth-based assays. (A) Schematic overview of the agar-based inhibition assay. 100 uL of an overnight culture of the *Streptococcus* isolates were spread on solid BHI agar plates and 20 µL of purified Lys_AF151 lysin (0,05mg/mL) were spotted on a sterile Whatman paper diffusion disk. Control spots included Cpl-1 lysin (0,05mg/mL) and elution buffer as negative controls. After 24 hours of incubation, inhibition zones were visually assessed and scored from IL = 0 (no inhibition) to IL = 3 (strong inhibition). (B) Representative results of broth-based growth curves of selected *S. salivarius* isolates in the presence (pink) or absence (blue) of the Lys_AF151 lysins (0,05 mg/mL). Growth was monitored via OD600nm every hour for 48 hours. Representative examples are shown for 4 *S. salivarius* isolates (AF23, AF187, AF316, AFB166) illustrating the range of inhibition observed. (C) Individual inhibition scores across four *Streptococcus* species: *S. salivarius* (N = 49), *S. parasanguinis* (N = 10), *S. mitis* (N = 9), and *S. thermophilus* (N = 2). Directionality arrows indicate agreement or divergence in inhibition level between agar and broth assays. Numeric values for IL are reported in Supp. Table 1 and Supp. Fig 2.

### Lys_AF151 lysin treatment shows specific activity against *S. salivarius* in stool-derived *in vitro* communities and is linked to higher *Coprococcus comes* relative abundance

To confirm the lytic activity and specificity of the Lys_AF151 lysin against *S. salivarius* in a complex community, we then evaluated its inhibitory potential in stool-derived *in vitro* communities (SIC). Half of each SIC generated where spiked with *S. salivarius* strains AF111ms or AF9ms, which were previously transformed with a plasmid that confers spectinomycin resistance (Table S1). The SICs spiked with *S. salivarius* and control SICs (no *S. salivarius* added) were treated with Lys_AF151 lysin at 0,05 mg/mL final concentration or the elution buffer as a negative control. CFU counting on selective media (BHI containing spectinomycin at 200 µg/µL) confirmed a significant decrease in *S. salivarius* abundance in lysin-treated SICs compared to untreated controls across all three SICs (Fig. 3A). This reduction was observed consistently for both *S. salivarius* strains tested (AF9ms and AF111ms), with an average decrease of 2–3 log units already apparent at the 24-hour time point. Notably, CFU counts for strain AF111ms were frequently lower than those for AF9 in lysin-treated samples, consistent with pure culture assays in which AF111ms exhibited a slightly higher susceptibility to lysin (Supp. Fig. 2). SICs were derived from three African donors, each displaying distinct bacterial community structures following stabilization in BHI medium (Fig. 3B&D).

**Figure 3.**
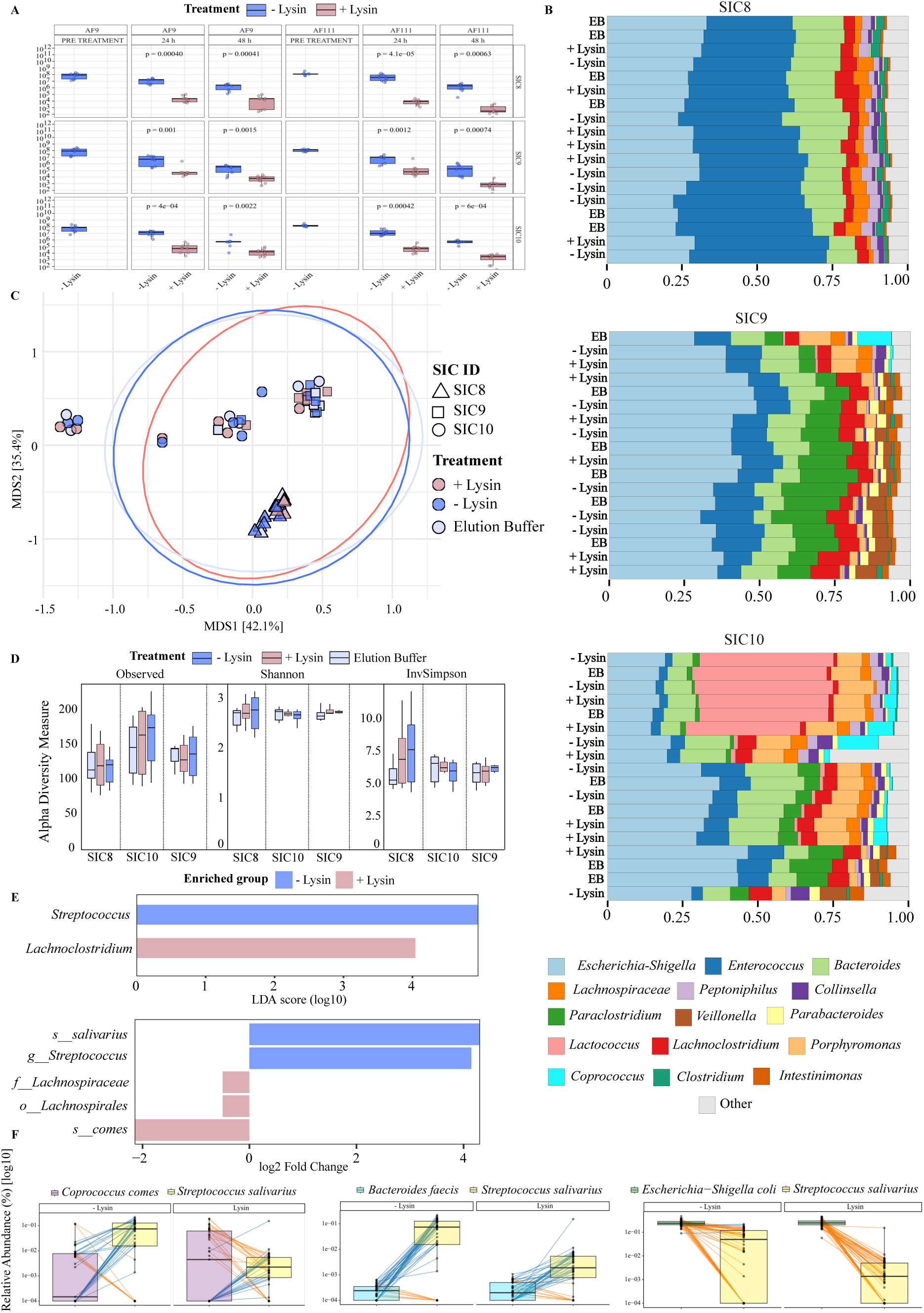
Lysin Lys_AF151 treatment modulates composition and abundance of stool-derived i*n vitro* microbial communities. (A) Quantification of *S. salivarius* colony-forming units (CFU/mL) over time across three SICs before and after lysin treatment. Boxplots show median and interquartile range, points are individual CFU measurements pooled across three SICs (SIC8A, SIC9A, SIC10A). Means ± SD are computed across measurements from the 3 SICs and comparison for + Lysin vs - Lysin was evaluated within each strain (AF9, AF111) and time (24, 48 h) using two-sided Wilcoxon rank-sum tests. p-values for each comparison are shown on the plot for each comparison. Blue = untreated controls; pink = lysin-treated. (B) Genus-level baseline relative abundance profiles of 16S rRNA gene amplicon sequencing data from the three stool-derived *in vitro* communities (SIC8, SIC9, and SIC10). All control SIC without spiked *S. salivarius* are shown (Untreated SIC; Lysin -, elution buffer-treated SIC; EB, and lysin-treated SIC; Lysin +). (C) Principal Coordinate Analysis (PCoA) of Bray-Curtis dissimilarities among the control SIC. (D) Alpha-diversity metrics (Observed, Shannon index and Inverse Simpson index) comparing untreated (green), elution buffer-treated (red) and lysin-treated (blue) SIC, without *S. salivarius* spiked. (E) Differential abundance analysis using Linear Discriminant Analysis (LDA) Effect Size (LEfSe). Upper panel: LDA scores showing taxa identified as significant biomarkers between treatment groups. Lower panel: relative abundances log2 fold change of key taxa differing between untreated (blue) and lysin-treated (pink) conditions. A negative log2 fold change indicates that the taxon is under-represented in the reference group (- Lysin) compared to the indicated condition. (F) Relative abundance trajectories of selected taxa from 16S data. Left, Middle and right: relative abundances of *S. salivarius* in pairwise comparison with other taxa (*Coprococcus comes, Bacteroides faecis, Escherichia-Shigella coli*).

As expected, lysin treatment of unspiked control SICs (i.e., without *S. salivarius* addition) had no measurable impact on overall community composition (Fig. 3C), and no taxa were identified as discriminatory biomarkers between treated and untreated samples by Linear Discriminant Analysis (LDA). In agreement with previous CFU counts, the 16S rRNA gene amplicon sequencing and LDA revealed that spiked SICs treated with lysin exhibited a significant reduction in the relative abundance of *S. salivarius* (Fig. 3E&F). For strain AF111ms, the mean relative abundance decreased from 11.69% ± 3.66% in untreated samples to 0.21% ± 0.13% following 24 h of lysin exposure (p < 0.0001, Wilcoxon rank-sum test), and from 8.79% ± 5.81% to 0.12% ± 0.05% after 48 h (p = 0.0002). Similarly, for strain AF9ms, the relative abundance decreased from 10.27% ± 7.48% in untreated controls to 0.66% ± 0.24% under lysin treatment after 24 h (p < 0.0001), and from 5.65% ± 6.04% to 0.46% ± 0.24% after 48 h (p = 0.0418). Interestingly, pairwise relative abundance analysis revealed a consistent increase in the relative abundance of *C. comes* in lysin-treated samples, which appeared inversely correlated with *S. salivarius* levels (Fig. 3E&F). We also performed this pairwise abundance comparison between *S. salivarius* and two common fecal commensals, *Bacteroides faecis* and *Escherichia coli*, which revealed no detectable correlation of *S. salivariu*s on the relative abundance of these two taxa (Fig. 3F). This analysis suggests a potential interaction of *S. salivarius* with *C. comes* and an absence of interaction with taxa such as *B. faecis* or *E. coli*.

### Lys_AF151 lysin treatment retains its specific activity against *S. salivarius in vivo* while promoting health-associated taxa

Next, to investigate the activity and potential use of the Lys_AF151 lysin *in vivo* under undernutrition conditions, we conducted experiments in a mouse model of childhood undernutrition^42^. Twelve 3-week-old undernourished C57BL/6J mice were colonized with *S. salivarius* AF111 by oral gavage on day 13 and 14 after arrival. On day 21, mice were divided into two groups (six cages, two mice per cage) and received either Lys_AF151 lysin (n = 6) or PBS as a control (n = 6) (Fig. 4A). Body weight and weight gain did not differ between treatment groups, indicating that the intervention was well tolerated (Fig. 4B & Supp Table 3).

**Figure 4.**
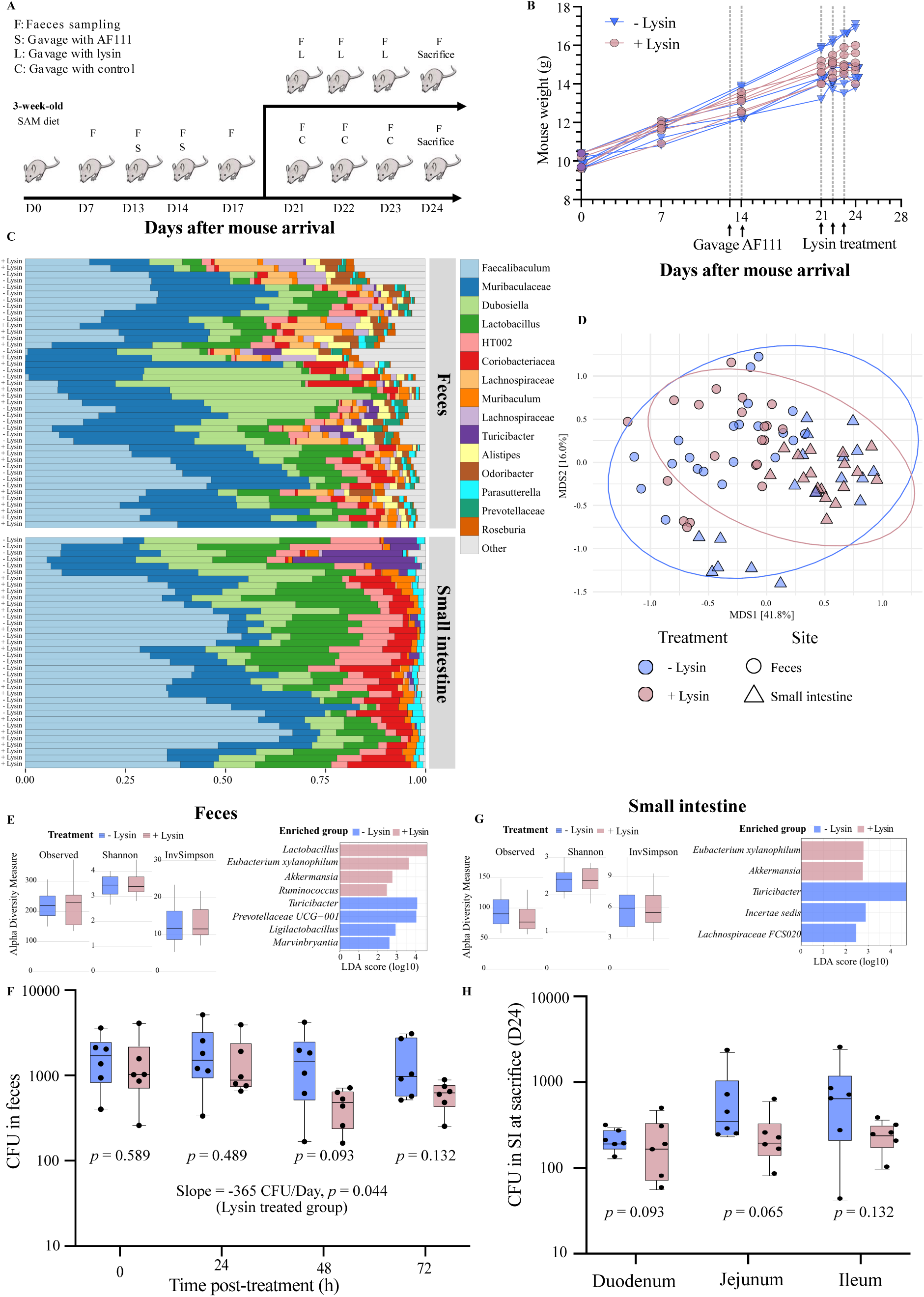
*In vivo* evaluation of lysin treatment in an undernourished mouse model of colonization by *S. salivarius*. (A) Schematic representation of the experimental design. Opportunist-pathogen free C57BL/6J male SPF mice (6 weeks old, SAM diet) were colonized with *S. salivarius* strain AF111 on day D13 and D14 post arrival through oral gavage. Mice were treated for 3 consecutive days with either the phage-derived lysin Lys_AF151 or vehicle control (PBS) by oral gavage on days D21 (T=0h), D22 (T=24h) and D23 (T=48h). Feces were collected at designated timepoints, and animals were sacrificed on day 24 (D24, T=72h) for analysis of small intestinal microbial community. (B) Longitudinal body weight measurements from day of arrival (D0) to sacrifice (D24). (C) Genus-level taxonomic composition of microbial communities in fecal (top) and small intestinal (bottom) samples from lysin and control-treated mice using 16S rRNA gene amplicon sequencing. (D) Principal coordinates analysis (PCoA) based on Bray–Curtis dissimilarity of genus-level relative abundances from fecal and small intestinal (SI) samples across treatment groups. Each point represents one sample (n = 78), colored by treatment (Lysin -, Lysin +) and shaped by site (feces vs. small intestine). (E) Effects of lysin treatment on microbial diversity and differentially abundant genera in fecal and (G) in small intestinal samples. Left: Alpha diversity metrics (Observed, Shannon diversity, and Inverse Simpson index) in fecal samples collected after lysin or control treatment. Right: Linear discriminant analysis effect size (LEfSe) showing genera differentially enriched between Lysin - and Lysin + groups. LEfSe was performed using CPM normalization at the genus level with Kruskal–Wallis and Wilcoxon p-value thresholds of 0.05 and an LDA score cutoff of 2.0, with multigroup stratification. (F) Quantification by qPCR of *S. salivarius* abundance in feces prior to the first treatment at day 21 and at 24 h (D22), 48 h (D23), and 72 h (D24) post-treatment. (H) Quantification by qPCR of *S. salivarius* abundance (CFU/mL) in the duodenum, jejunum, and ileum (H) after sacrifice at 72 h post-treatment. Bars represent mean ± SD. CFU: colony forming units; PLT: post-lysine treatment.

*S. salivarius* loads were quantified by qPCR over time in feces and at endpoint across intestinal compartments. At D21 (treatment baseline, T=0 h), fecal counts were comparable between groups: 1,472 ± 1,351 CFU/mL (Lysin) vs 1,745 ± 1,120 CFU/mL (Control), *p* = 0.589. At D22 (24 h after first dose), values remained similar: 1,500 ± 1,268 CFU/mL (Lysin) vs 2,031 ± 1,699 CFU/mL (Control), *p* = 0.485). By D23 (48 h after dose 1), we started to observe lower counts with lysin relative to control: 456 ± 215 CFU/mL (Lysin) vs 1,641 ± 1,432 CFU/mL (Control), *p* = 0.093), corresponding to a Control/Lysin-treated fold-change of ≍ 3.60. This pattern persisted at D24 (72 h after dose 1), with 603 ± 219 CFU/mL (Lysin) vs 1,467 ± 1,121 CFU/mL (Control), *p* = 0.132) with a Control/Lysin fold-change of ≍ 2.43.

Although previous day-wise comparisons did not cross the *p* < 0.05 threshold, the time-dependent fold-change profile suggests a treatment-associated reduction that becomes apparent from 48 h onward (Fig. 4F). A trend analysis performed with a linear regression model showed a significant negative trend in mice fecal CFU counts over time for the Lysin group (Slope = −365.2, *p* = 0.044), whereas no trend was detected in PBS treated controls (Slope = −122.7, *p* = 0.613). In small intestinal tissues, lysin effect was not observable in the duodenum : 203 ± 154 CFU/mL (Lysin) vs 208 ± 62 CFU/mL (Control), *p* = 0.485. However, lower burdens became apparent for the lysin treated mice in the jejunum : 244 ± 181 CFU/mL (Lysin) vs 675 ± 773 CFU/mL (Control), ≈2.8-fold reduction, *p* = 0.065. Similar results were observed in the ileum: 237 ± 91 CFU/mL (Lysin) vs 795 ± 834 CFU/mL (Control), ≈3.3-fold reduction, *p* = 0.132; Fig. 4H & Supp Table 3). Notably, variability was high in controls for jejunum/ileum, whereas the lysin groups showed lower SDs, consistent with a more uniformly reduced *S. salivarius* burden in the distal small intestine. These findings are consistent with our results obtained *in vitro* and demonstrate that orally delivered lysin reaches the intestinal lumen and reduces target bacterial load across compartments.

Microbial community analysis by 16S rRNA gene amplicon sequencing revealed minimal disruption to the overall microbiota *in vivo* as demonstrated in a principal coordinates analysis (PCoA) based on Bray-Curtis distances (Fig. 4C). As expected, we observed a strong separation between fecal and small intestinal (SI) communities, consistent with gut biogeography (Figure 4C). Alpha diversity metrics (Observed, Shannon, and Inverse Simpson indices) remained stable across groups, indicating that the treatment did not affect overall microbial richness or evenness in either feces or SI (Fig. 4E,G). Detailed compositional analysis revealed few noteworthy treatment-associated effects. In feces, linear discriminant analysis identified relative enrichment of *Lactobacillus, Eubacterium xylanophilum* group*, Akkermansia* and *Ruminococcus* in lysin-treated animals, while *Turicibacter, Ligilactobacillus, Prevotellaceae UCG-001 and Marvinbryantia* were found to be higher in samples from untreated mice (Fig. 4E). In the SI samples, as observed in the feces, lysin-treated animals showed a significant enrichment in the *Eubacterium xylanophilum* group and *Akkermansia*, while *Turicibacter* was more abundant in the control group (Fig. 4F). The *Lachnospiraceae FCS020* group and *Incertae sedis taxa* were enriched in controls compared to lysin-treated ones, but only in the SI.

### *S. salivarius* exhibit niche overlap with numerous taxa involved in the production of butyrate

Given the *in vitro* decrease in *S. salivarius* alongside increases in butyrate producers (*C. comes* and *E. xylanophilum* group) after lysin treatment, we hypothesized that *S. salivarius* may compete with these taxa, potentially explaining the lysin’s indirect effects. To investigate this hypothesis, we conducted spent media assays^43^. These assays were performed on a subset of 10 *S. salivarius* “target” strains, representing the different genomic clusters (C1-C4) of the *S. salivarius* diversity based on whole genome average nucleotide identity (ANI) from our *S. salivarius* isolate collection (Fig. 5B). These strains were then cultured in the spent medium of a corresponding “donor” strain. Growth performance in spent medium was assessed relative to that in fresh BHI medium (Fig. 5A), with reductions in growth rate in spent medium indicating that both strains likely exploit similar nutrient resources, consistent with metabolic niche overlap. When *S. salivarius* was cultured in the *C. comes* spent medium, we observed delayed and reduced growth relative to growth in fresh BHI, concordant with our hypothesis based on the negative interaction observed in the SICs (Fig. 5C and Fig. 3F). To expand this analysis, we assessed the growth of 10 additional *S. salivarius* strains in the spent medium of 33 different species of bacteria involved in butyrate production (Supp. Fig. 3, also (Supplementary Table 2). For the 10 *S. salivarius* strains tested (Supplementary Table 1), growth was almost completely abolished in 10 out of the 33 spent media and impaired in 19 out of 33, thus suggesting potential niche overlap with 29 of the 33 (88%) butyrate-producers tested (Supp. Fig. 3, Supp. Table 2). In cases of partial impaired growth, strain-specific differences among *S. salivarius* isolates were obvious, indicating that strength of competitive outcomes may vary depending on the specific pairings of “donors” and “target” strains involved. Only four donor-derived spent media did not appear to inhibit *S. salivarius* growth relative to fresh BHI, suggesting considerable niche separation. Tested species that did not show overlap with *S. salivarius* included one strain of *B. faecis* (EBT048), two strains of *Anaerobutyricum hallii* (EBT072, EBT151), and one strain of *Coprococcus catus* (ATCC27761). The absence of *S. salivarius* growth defect in *B. faecis* spent medium and the absence of co-variation between these two species observed in the previous *in vitro* 16s rRNA community analysis (Fig. 3F) further support the experimental approach for identifying competitive (or neutral) interactions within the microbiota.

**Figure 5.**
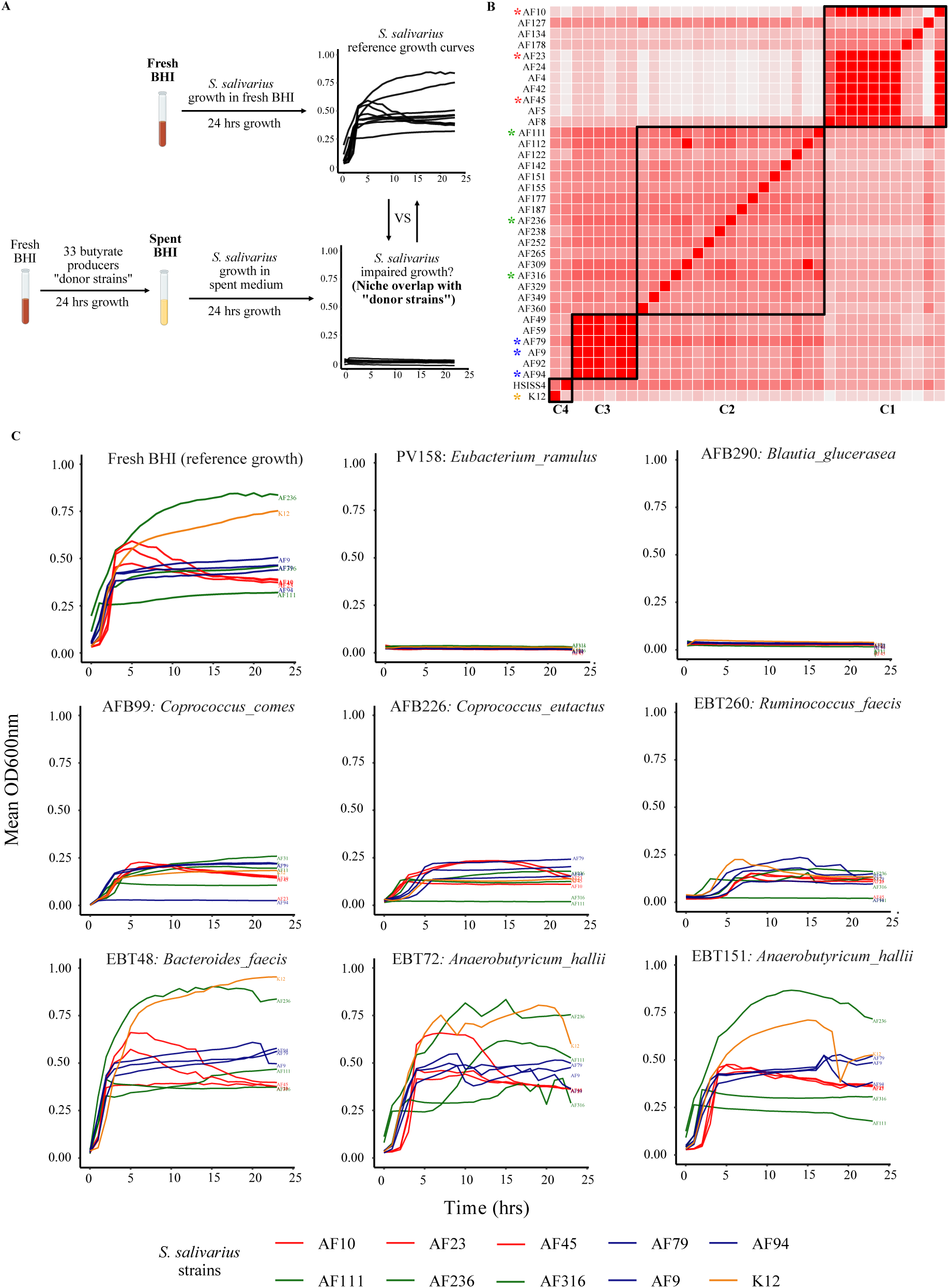
Growth inhibition of *S. salivarius* by spent media from different bacterial species. (A) Schematic representation of the spent medium assay. Fresh BHI and spent media (spent BHI) were used to culture the 10 “target” *S. salivarius* strains. Spent medium was produced by growing 33 “donor” strains (listed in Supp. Table 2) in fresh BHI to produce spent medium (spent BHI). (B) The 10 “target” *S. salivarius* strains used for spent medium assays were selected to represent different genomic clusters (C1-C4), based on whole genome average nucleotide identity (ANI) from our *S. salivarius* isolate collection (listed in Supp Table 1). Darker red indicates higher sequence identity between genomes, with a maximum of 100%, while lighter shades represent lower identity, down to a threshold of 70 % (absolute white). (C) Selected examples depicting the growth of 10 *S. salivarius* strains in spent medium from different “donors”. Colors for growth curves were assigned to *S. salivarius* strains according to the ANI cluster they belong (Red : cluster 1 = C1, Green : cluster 2 = C2, Blue : cluster 3 = C3, Orange : cluster 4 = C4).

## DISCUSSION

Childhood stunting remains a major global health challenge, with mounting evidence pointing to the contribution of microbial dysbiosis in its aetiology. In particular, small intestinal oral bacterial overgrowth (SIOBO), characterized by ectopic colonization by oral taxa such as *S. salivarius*, has been associated with persistent inflammation, impaired nutrient absorption, and disruption of beneficial taxa including butyrate producers^3,7,12,13^. This aberrant microbial configuration does not appear to be reversible by traditional dietary interventions alone^44^, underscoring the need for adjunct therapeutic strategies. Precision microbiota-targeted interventions offer a promising approach to address dysbiosis as part of a multifactorial strategy to promote intestinal recovery and functional nutrient absorption in undernourished children and other gut related diseases^16,23^.

In this study, we investigated the feasibility of using phage lysins to control *S. salivarius* in the complex gut environment. To achieve this, we first screened our collection of *S. salivarius* isolates, which identified two complete prophage sequences in strains AF151 and AF238. Each prophage genome harbors a single gene encoding for an endolysin (Lys_AF151 and Lys_AF238), both containing a hydrolase catalytic domain and cell wall and choline-binding domains, which are characteristic of streptococcal phage endolysins^34,45^. Our successful attempt to induce temperate phages from both strains, with genomes 100% similar to the prophage sequences, confirmed functionality of both prophages and supported Lys_AF151 and Lys_AF238 as functional cell wall hydrolases to release phage progeny. Following cloning and overexpression in *E. coli*, we purified Lys_AF151 for detailed functional characterisation.

Our host range assays demonstrated potent and specific lytic activity of Lys_AF151 against *S. salivarius*, with negligible activity against closely related *Streptococcus* species, including *S. parasanguinis*, *S. mitis*, and *S. thermophilus*. This specificity is critical for applications aiming to rationally modulate the microbiota, avoiding unsought detrimental off-target disruption of commensals. Strain-specific variability was noted between tested isolates, such as AF111 showing consistently higher susceptibility than AF9 across both agar and broth assays, a pattern also confirmed in our SIC experiments. These differences may be due to variations in surface-accessible peptidoglycan motifs or differential susceptibility to cell wall degradation^46,47^ and highlight the importance of considering inter-strain structural variation when designing lysin-based interventions. Future work could explore whether variations in lysin activities correlate with specific bacterial genomic traits related to cell wall composition or surface protein expression and could focus on how to obtain a more robust lytic activity for a larger diversity of specific target strains. We also observed minor inter-SIC variability in lysin efficacy, which may reflect protective influences from environmental factors such as pH fluctuations, ionic strength, redox potential, or the presence of bile salts^23,48^. Additionally, features of the microbial community itself, such as the presence of lysin-sequestering species, lysin-inactivating proteases or biofilm formation, may further cause variations in lysin accessibility or activity. These complex interactions warrant deeper mechanistic investigation to better understand the determinants of lysin performance in diverse microbial and environmental contexts. Nevertheless, we observed a robust reduction of *S. salivarius* across multiple settings, including pure cultures, complex communities, and an *in vivo* mouse model, which is encouraging and support further exploration of lysins in both mechanistic studies and potential clinical applications.

The SIC model is a valuable tool for preclinical microbiome research, as it preserves complexity while enabling controlled perturbations^49^. When applied to human stool-derived *in vitro* communities, Lys_AF151 selectively reduced spiked *S. salivarius* strains without perturbing overall community structure or diversity. *In vivo*, oral administration of the Lys_AF151 lysin also reduced *S. salivarius* loads in mice, without inducing observable adverse effects or significant changes in host body weight or intestinal microbiota composition. The ∼2-log reduction observed in both fecal and small intestinal *S. salivarius* reflects the effects seen in SICs, further supporting the translational relevance of the SIC model. Community profiling by 16S rRNA gene amplicon sequencing confirmed the targeted effect of Lys_AF151 and identified a potential co-exclusion between *S. salivarius* and *C. comes*. *C. comes* is a known butyrate-producing bacterium, and its expansion likely reflects ecological release following the removal of a competitor^50^. Similarly, *in vivo* analysis highlighted the enrichment of bacterial taxa broadly associated with beneficial health outcomes, including *Lactobacillus*, *Akkermansia* and members of the *E. xylanophilum* group, the latter also known for its butyrate-producing capability. *Lactobacillus* species are well-characterized for their probiotic properties, contributing to barrier function, acidification of the gut environment, and competitive exclusion of pathogens^51^. Similarly, *A. muciniphila*, a mucin-degrading bacterium residing in the intestinal mucus layer, has been linked to improved metabolic health and reduced inflammation in both preclinical and clinical studies^52–54^. The *E. xylanophilum* group is known for its capacity to ferment xylan to produce formate, acetate and butyrate^55^. *Ruminococcus* species like *R. bromii* are central to resistant starch degradation and support butyrate-producing cross-feeders^56^. These compositional shifts might reflect indirect effects of *S. salivarius* removal on the host microbial ecosystem. These likely include indirect metabolic interactions within the ecosystem such as altered nutrient availability or niche competition. It is noteworthy that butyrate-producing taxa were enriched under lysin-treated conditions both *in vitro* and *in vivo, with C. comes* increasing in stool-derived *in vitro* communities and *E. xylanophilum* in the mouse model. This negative correlation between *S. salivarius* and butyrate producers is particularly interesting, as similar negative correlations has been previously observed in a study on childhood undernutrition^3^ and intestinal bowel disease (IBD)^57^.

These observations led us to further investigate the interactions between *S. salivarius* and other butyrate producing species. We performed spent medium assays and showed that all the 10 *S. salivarius* strains tested exhibited impaired growth in the spent media from 29 of 33 butyrate-producing isolates, including *C. comes*. This widespread growth inhibition suggests extensive metabolic niche overlap and competition for shared resources, potentially explaining the inverse correlation between *S. salivarius* and *C. comes* observed in our experiments. Furthermore, our data might explain the co-exclusion patterns seen in sequencing data from studies on undernutrition and IBD^3,57^. We also stratified *S. salivarius* isolates into genomic clusters using whole genome average nucleotide identity (ANI) and observed that strains within the same cluster often exhibited similar inhibition patterns across different spent media. Similar inhibition patterns were observed among strains within the same cluster, particularly in cases of partial growth inhibition in spent media. This observation supports the notion that genomic relatedness may reflect metabolic similarity^58^ and further highlights strain-specific interactions as key determinants of bacterial dynamics in the gut environment^59,60^. Future studies could investigate whether lysin-mediated targeting of specific bacterial taxa systematically promotes the growth of other microbes occupying overlapping ecological niches.

While our *in vivo* experiments and analyses primarily focused on the reduction of *S. salivarius* bacterial load, future studies could investigate whether such targeted depletion leads to improvements in physiological markers of gut health, including intestinal barrier function, reduced inflammation, enhanced nutrient absorption, and increased production of health-associated metabolites such as butyrate and other short-chain fatty acids. This will be particularly relevant considering the observed indirect enrichment of butyrate-producing taxa following lysin treatment. Moreover, a better understanding of lysin pharmacokinetics, proteolytic stability, and optimal delivery formulations will be critical to enhancing and sustaining efficacy *in vivo*. Last, it will be important to assess for long-term stability of the new ecosystem established post lysin treatment and its overall long-term impact on global host physiology.

To date, only a limited number of *in vivo* studies have investigated the effects of lysins on the intestinal microbiome. Harhala *et al.* reported that intraperitoneal administration of the pneumococcal lysins Pal or Cpl-1 did not result in significant alterations to the microbiome composition in mice^61,62^. In contrast, administration of an enterococcal lysin LysEF-P10 led to observable, though not statistically significant, changes in the microbial community^62^. Our results are thus in line with these previous studies underlining low off-target effects and the possible precision engineering of the microbiome using phage lysins.

Importantly, our findings also highlight the utility of lysins not only as therapeutic agents but also as precision tools to probe ecological relationships. Unlike antibiotics or broader-spectrum antimicrobials, lysins enable species-level perturbation within complex communities, offering a unique opportunity to study causal interactions. In this study, the depletion of *S. salivarius* revealed its negative interaction with multiple taxa, suggesting niche overlap and potential ecological release of competing species after *S. salivarius* reduction^63^.

The ability to design and apply additional lysins targeting specific taxa of interest could allow us to test ecological hypotheses experimentally and help in disentangling causality in bacterial networks that may underlie dysbiosis or gut related diseases.

Moreover, the identification of functional lysins within prophages offers practical advantages. Unlike lytic phages, which can be difficult to find, isolate and propagate, prophage-encoded enzymes are accessible from sequenced bacterial genomes and amenable to synthetic production. With larger bacterial genome databases and advances in DNA synthetic biology^64,65^, lysins can now be more easily designed and optimized from sequence data alone, without requiring the isolation of either host or phage. This opens the door to rational design of lysin cocktails targeting combinations of pathobionts or keystone taxa to explore ecological relationships.

Altogether, this work demonstrates that phage-derived lysins can serve as precise, species-specific tools to modulate microbial communities. Beyond therapeutic use, lysins offer a valuable means of probing microbial ecology by enabling targeted, temporal removal of taxa to infer interactions, dependencies, and emergent community behaviour. In the context of SIOBO and childhood undernutrition, our findings provide a proof-of-concept and a foundation for microbiota-based interventions that extend beyond specific bacterial depletion. It also offers mechanistic insights into how specific taxa modulate health-relevant microbial networks. Seen that oral bacteria are also implicated in many other inflammatory and non-inflammatory diseases, Lys_AF151 might further be of interest to treat other ailments linked to *S. salivarius* overgrowth^2^.

By combining microbial precision targeting with ecological investigation, lysin-based strategies present a dual opportunity: to both treat and better understand dysbiosis-driven diseases. As we refine these tools and expand their applications, lysins might become invaluable in the broader effort to design microbiota-targeted and informed interventions for global health.

## MATERIALS AND METHODS

### Ethics statement

The samples and strains used in this study stem from the Afribiota project^66^, which was initially approved by the Institutional Review Board of the Institut Pasteur (2016-06/IRB) and the National Ethical Review Boards of Madagascar (55/MSANP/CE, 19 May 2015). All participants received oral and written information about the study. The legal representatives of the children provided written consent to participate in the study and for data re-use. The research was performed in accordance with all relevant guidelines and regulations and was performed in accordance with the Declaration of Helsinki. The present analysis (AfribiotaScreen) was approved by the Swiss National Ethics board under number (BASEC-ID 2020-03058).

### Isolation of Streptococcus salivarius Strains

*Streptococcus salivarius* strains used in this study (Supp Table 1) were isolated from fecal samples collected from African child donors. Frozen fecal material was processed under anaerobic conditions by diluting in pre-reduced phosphate-buffered saline (PBS) and plating onto Brain Heart Infusion (BHI) agar (BD Difco, Cat. No. 279830, Franklin Lakes, NJ, USA) supplemented with 1 g/L inulin from chicory (Sigma-Aldrich, Cat. No. I2255, Darmstadt, Germany). Plates were incubated in a strictly anaerobic workstation (Coy Laboratories, Ann Arbor, MI, USA) under a gas mixture of 7% H₂, 20% CO₂, and 73% N₂ at 37 °C for 3–4 days. Individual colonies were selected and purified through three rounds of restreaking on fresh BHI-inulin agar. Species identification was performed using MALDI-TOF MS^67^, and verified isolates were stored in 20% glycerol at −80 °C until further use.

### Prophage Induction Assay

To perform prophage induction, 40 *S. salivarius* strains were cultured with three concentrations of mitomycin C (0.156 µg/µL, 0.078 µg/µL, or 0.039 µg/µL) (Sigma-Aldrich, REF 50-07-7). Overnight pre-cultures were prepared by inoculating each strain into 1.5 mL of BHI broth and incubating at 37 °C overnight. The following day, 360 µL of each overnight culture was transferred into 8.64 mL of fresh BHI in culture tubes and incubated at 37 °C for 1 h. After this incubation, mitomycin C solution was added to each culture at one of three final concentrations. Bacterial growth was monitored in parallel in a 96-well plate, using a Synergy H1 microplate reader (BioTek) to follow optical density at 600 nm (OD600) over 24 h. Following incubation, cultures were centrifuged, and the supernatant was filtered through 0.22 µm filters (ClearLine, #257160) and stored at 4 °C. For phage concentration, 2 mL of filtered supernatant was centrifuged at 20,000 × g for 16 h at 4 °C. The resulting supernatant was discarded, and the phage-containing pellet was resuspended in PBS and stored at 4 °C for downstream applications.

### Electron microscopy

To prepare the samples for electron microscopy, 0.5 mL of the induced supernatants were centrifuged for 15 minutes at 23 °C and at 8000 x g in an Amicon Ultra-0.5 mL centrifugal filter (Merck, Germany, REF C82301) to concentrate the potential phages. The sample were prepared by negative staining with uranyl acetate, and imaging of the samples were performed by the EM Facility at the University of Lausanne.

### Bacterial and phage DNA extraction

Genomic DNA from single strains or SIC communities was extracted using the Maxwell RSC PureFood GMO and Authentication Kit (Promega, REF AS1600), following the manufacturer’s protocol with slight modifications. Briefly, 200 µL of bacterial culture/SIC was added directly to 800 µL of CTAB buffer in a PowerBead Pro Tube (Qiagen, REF 19301). Lysis was performed using a vortex bead-beater (Scientific Industries, REF SI-0276) at maximum speed (level 10) for 10 min at room temperature. After centrifugation at 14,000×g for 10min, 600 µL of the supernatant was transferred to a 1.5mL Eppendorf tube. To degrade protein and RNA, 40 µL of Proteinase K and 20µL of RNase A (Promega, REF AS1600) were added and the mixture was vortexed briefly and incubated at 70°C for 10 min. DNA clean-up was completed using the Maxwell RSC 48 instrument (Promega, REF AS8500) according to the PureFood protocol. Phage DNA was extracted following prophage induction with mitomycin C (Sigma-Aldrich, REF 50-07-7). A total of 2 mL of culture supernatant was collected and centrifuged at 20,000×g for 16h at 4°C. The resulting phage pellet was resuspended in 490 µL of PBS. To remove residual bacterial DNA, 10µL of DNase and 70µL of Buffer RDD (RNase-Free DNase Set, Qiagen, REF 79254) were added and the sample incubated in a water bath at 37°C for 1h. DNase was inactivated by heating at 75°C for 10 min. Next, 500 µL of the treated suspension was transferred to a PowerBead Pro Tube and combined with 500 µL of CTAB buffer (Promega, REF AS1600). Capsids were lysed using a vortex bead-beater (Scientific Industries, REF SI-0276) at maximum speed for 5min at room temperature. The lysate was centrifuged at 14,000×g for 10 min, and 600 µL of the supernatant was transferred to a 1.5mL Eppendorf tube. Protein and RNA were degraded by adding 40 µL of Proteinase K and 20 µL of RNase A (Promega, REF AS1600), followed by vortexing and incubation at 70°C for 10 min. Final purification of phage DNA was completed using the Maxwell RSC 48 instrument (Promega, REF AS8500) and the PureFood protocol.

### Bacterial and phage whole-genome sequencing

Whole-genome sequencing of bacterial and phage genomic DNA was performed by Novogene UK (Cambridge, UK). Libraries were prepared using a microbial whole-genome library preparation kit with an average insert size of 350 bp. Sequencing was carried out on the Illumina platform using paired-end 150 bp (PE150) reads, generating approximately 1 Gb of raw data per sample. Raw FASTQ reads were processed using Fastp^68^ with default parameters for quality filtering and adapter trimming. Cleaned paired-end reads were assembled using SPAdes^69^ in paired-end mode with default settings. Genome annotation was performed using Prokka^70^ with default parameters. Filtered reads were aligned to the reference bacterial genome using Bowtie2^71^. Termini and packaging mechanisms were predicted using PhageTermVirome^40^ with default settings. Comparative analysis of phage-encoded proteins was carried out using Clinker^72^. Lysin sequences were identified within assembled phage genomes based on functional annotations and were further analyzed for conserved domains using InterProScan^73^.

### 16s rRNA gene amplicon sequencing using primers targeting the V3-V4 regions

Bacterial community composition was assessed by 16S rRNA gene amplicon sequencing. The V3-V4 region of the 16S rRNA gene was amplified using primers 341F/806R, from DNA previously extracted from SIC cultures or mouse feces. Amplicons were generated and sequenced at Novogene UK (Cambridge, UK), on an Illumina platform using paired-end 250 bp reads. A minimum of 30k sequence tags per samples were obtained. Raw sequences were processed in R using the DADA2^74^ pipeline, which includes quality filtering, dereplication, denoising, merging of paired-end reads and removal of chimeric sequences to generate amplicon sequence variants (ASVs). Taxonomic assignment of ASVs was performed using the latest SILVA database^75^, as of July 1st 2025 (silva_nr99_v138.1_wSpecies_train_set.fa.gz). Downstream analyses, including diversity metrics, community composition, and statistical testing were conducted using the phyloseq package^76^.

### Cloning of the Lys_AF151 lysin

The coding sequence for the lysin Lys_AF151 was inserted into a pET-based bacterial expression vector together with a C-terminal polyhistidine tag. The resulting plasmid was transformed into BL21(DE3) Gold *E. coli* cells, and a 1 L culture was grown in TB medium to an O.D. 600nm of 1.0 at 37°C. The temperature was then reduced to 18°C and expression induced by adding IPTG to a final concentration of 0.4 mM. After an overnight (16 h) incubation, the cells were harvested by centrifugation and the pellet was resuspended in 70 ml of Lysis Buffer (50 mM Tris-HCl pH7.5, 300 mM NaCl, 5 % glycerol, 25 mM Imidazole) containing protease inhibitor cocktail (Sigma) and 5 mM beta-mercaptoethanol (Sigma). Cells were lysed by sonication (Bandelin sonicator, 40 % output, 10 min, 1 sec ON / 1 sec OFF) on ice, and the lysate was clarified by centrifugation at 40000 x g for 30 minutes at 4°C. The clarified lysate was loaded onto a 5 ml HisTrap-HP column (Cytiva) and the column was then washed with 5 column volumes (cV) of Lysis Buffer. Bound proteins were eluted with a 10 cV gradient from Lysis Buffer to Lysis Buffer supplemented with 500 mM Imidazole. Peak fractions were pooled and either used directly for *in vitro* experiments or dialyzed overnight (16 h) into PBS for *in vivo* experiments on mice.

### Lysin inhibition assay

To evaluate the inhibitory activity of the phage-derived lysin Lys_AF151, a host range assay was performed against 49 clinical isolates of *S. salivarius*. Each strain was cultured overnight in 3 mL of BHI broth at 37 °C. The following day, 100 µL of each overnight culture was plated onto BHI agar plates using sterile glass beads. Sterile Whatman paper disks were manually prepared using a hole punch and were placed on inoculated agar plates. Then, 20 µL of each solution of Lys_AF151, Cpl-1, or the elution buffer (EB) was pipetted onto individual disks. The EB, containing190 mM imidazole, served as a negative control for the *in vitro* experiments, as it was used for lysin elution during protein purification. Plates were incubated at 37 °C overnight. Zones of inhibition were visually assessed based on the clarity of the lysis area and classified into four distinct inhibition levels (IL): IL0: No inhibition, IL1: Low inhibition, IL2: Moderate inhibition and IL3: High inhibition. For inhibition assays in liquid culture, all tested strains were first grown overnight in BHI medium. Pre-cultures were then restarted by inoculating 8 µL of overnight culture into 172 µL of fresh BHI in 96-well plates. Lysin was added at a final concentration of 0.05 mg/mL by pipetting 20 µL of a 0.5 mg/mL lysin stock solution (or 20 µL of BHI or elution buffer controls) into each well, bringing the final volume to 200 µL. All conditions were tested in triplicate.

### Generation and stabilization of stool-derived *in vitro* communities (SICs)

SICs were derived from three distinct stool samples collected from children in Madagascar. SIC8 and SIC10 originated from fecal samples of severely malnourished children, while SIC9 was derived from a normally nourished child. The creation of the SICs was based on the protocol from Aranda-Díaz *et al.*^49^. Briefly, to establish the SICs, 0.2 g of frozen stool from African children was resuspended in 3 mL of pre-reduced phosphate-buffered saline (PBS) inside an anaerobic chamber containing a gas mixture of 7% H₂, 20% CO₂, and 73% N₂.

Samples were left at room temperature for 10 minutes to allow hydration of larger stool fragments, followed by vortexing for 2 minutes and pipetting to ensure homogenization. After settling for 1 minute at room temperature, 50 µL of the supernatant was transferred into 10 mL of pre-reduced BHI medium (Difco Brain Heart Infusion, Becton Dickinson, REF 237500) supplemented with 1 g/L inulin from chicory (Sigma-Aldrich, Cat. No. I2255). Each sample was inoculated in triplicate and incubated at 37 °C under anaerobic conditions. SICs were passaged every 48 hours by transferring 5 µL of vortexed culture into 1 mL of fresh BHI-inulin medium in 96-deep-well culture plates (Nunc 96 DeepWell plate, non-treated, Merck, Cat. No. Z717274-60EA) at a 1:200 dilution. Plates were covered with lids to minimize evaporation. A total of five passages were performed to stabilize the communities. After the final passage, 500 µL of each SIC was mixed with 500 µL of pre-reduced 40% glycerol containing palladium beads (Precipitated palladium, Sigma-Aldrich, Cat. No. 520810) and stored at −80 °C until further use. For revival, frozen SICs were inoculated into 1 mL of fresh BHI-inulin medium and incubated for 48 hours under anaerobic conditions, followed by one additional passage to re-stabilize the communities.

### Assessment of the Lys_AF151 lysin activity in stool-derived *in vitro* communities

To assess the bactericidal effect of the lysin Lys_AF151 against *S. salivarius* in complex bacterial community, we tested it against two strains of *S. salivarius* (AF9ms and AF111ms) in the three different stool derived *in vitro* communities (SICs) previously created and stabilized. All experiments were conducted under anaerobic conditions. Initially, SICs were grown from glycerol for 48h and pure bacterial cultures for 24h in BHI at 37 °C. Following the overnight incubation, 80 µL of the precultured strains and 40 µL of the SICs were mixed in 1 mL of fresh BHI in a 96 deep well plate and incubated for 24h at 37°C. Before adding the first lysin treatment, aliquots of the different conditions were plated for CFU counting and a part of the sample was collected for future sequencing analysis. The first dose of lysin Lys_AF151 with a final concentration of 0.05 mg/mL was added to the mix of *S. salivarius* and SIC. For the control we added either BHI or the elution buffer. After 24 h the different conditions were plated for CFU counting, samples were collected for future sequencing analysis and the second dose of lysin Lys_AF151 was added (0.05 mg/mL). After 24 h, the different conditions were again plated for CFU counting and samples were collected for future sequencing analysis.

### Quantification of S. *salivarius* by qPCR

The total abundance of *Streptococcus salivarius* was assessed using strain-specific primers designed using the PUPppy primer design pipeline^77^ and the SYBR Green Universal Master Mix (Thermo Fisher) following manufacturer’s protocol. Primers were used at a final concentration of 0,370 uM, in a final reaction volume of 15 uL. Sequences of the qPCR primers for strain AF111 quantification are TCGTTGTCAGGAGTGGGTTG (AF111_Forward) and TCTTATCCGACCAGCCTTGC (AF111_Reverse). The concentration of extracted DNA was measured by Nanodrop and subsequently diluted to 20 ng/μL. qPCR was performed for each sample in triplicate using 384-well plates and a QuantStudio 5 Real-Time PCR Machine. Ct values were interpolated to colony-forming unit (CFU) using a standard curve generated from qPCR amplification of purified *S. salivarius* AF111 DNA. Results were expressed in CFU/mL of culture.

### Animal experiments

All animal experiments were ethically approved by the responsible authorities (Directorate General for Agriculture, Viticulture and Veterinary Affairs, Cantonal Veterinary Office Vaud, License VD3965b). Specific- and opportunist-pathogen free (SOPF) male C57BL/6JRj mice (3-week-old) were purchased from Janvier-labs (France). Mice were maintained on a 12-h light/dark cycle (07:00-19:00) in a temperature- and humidity-controlled environment (23 ± 2 °C and 55% ± 10%, respectively). Food and water were provided *ab libitum*, with food consumption monitored to avoid overeating. All mice were kept under specific pathogen-free (SPF) conditions (autoclaved cages, bedding, water bottles and water) in the SPF animal facility at the University of Lausanne, Switzerland. Upon arrival the animal facility, all mice were randomly split into experimental cages (two mice per cage, three cages per condition) and placed on a Severe Acute Malnutrition (SAM) diet based on a porcine model of infant malnutrition^42^ (MAIS +1800 mg/kg Sodium, SAFE R8960A01R 00128; Safe, France). The SAM diet composition is as follow: 322.5 kcal/kg (9.2%) from protein, 331.5 kcal/kg (9.5%) from fat, and 2,835 kcal/kg (81.3%) from carbohydrates, totaling 3,489 kcal per kilogram. *S. salivarius* AF111 was cultivated overnight at 37 °C in TSB medium, resuspended in 1x PBS and 5×10^8^ CFU/mL were introduced to all the mice via oral gavage on days 13 and 14 after arrival. Seven days post-colonization (day 21), 100 uL of lysin solution (400 ug total) in 1x PBS or 1x PBS alone as a control was administered by oral gavage for three consecutive days. Mice were transferred to fresh cages after the first treatment to minimize gut recolonization from the environment. Weight of the mice was followed, and feces were collected during the lysin or placebo intervention until day 24 post-arrival, when mice were euthanized. Bacterial CFU counts were determined from fecal samples collected on four consecutive days (D21–D24) as well as from intestinal segments (duodenum, jejunum, and ileum) at sacrifice. For fecal samples, each time point included six mice per treatment group and organs were sampled at sacrifice from the same animals (n = 6 per group). For each day and organ, values are reported as individual measurements together with group means ± standard deviation (SD) (Supp Table 3). Body weights were recorded longitudinally for each mouse at arrival (Day 0) and subsequently at Day 7, Day 14, and Days 21–24. Weight gain relative to baseline (Day 0) was calculated for each mouse at Day 21–24, and raw values are presented along with group means ± SD (Supp Table 3). Statistical analyses were performed in R (version 2024.12.1+563) using the packages dplyr, tibble, tidyr, and broom. To compare Lysin-treated and control groups at each time point for fecal CFU, organ CFU, body weight, and body weight gain, we applied the Wilcoxon rank-sum test (two-sided Mann–Whitney U test). To evaluate temporal changes in fecal CFU counts across days, replicate-level data were modeled using linear regression (CFU ∼ day) separately for each group, with regression slopes (CFU/day), standard errors (SE), and p-values reported (Supp. Table 3).

## DATA AVAILABILITY STATEMENT

Genome and associated sequencing reads will be released on the ENA and SRA server upon publication. Request regarding strains should be directed to the corresponding author Pascale Vonaesch.

## AUTHOR CONTRIBUTIONS

PV, JRG and GR initiated the study. JRG, GR and PV supervised the work. PV, JRG and AA designed the study. JRG, AA, YT, XW, SM, PG, SY and MT performed the experiments. GR, CHC and JM provided critical input on specific technical aspects of the project. JRG, AA, YT and XW performed the data analysis. JRG and PV wrote the initial draft. All co-authors revised the final draft.

## ACKNOWLEDGEMENT

The authors would like to thank the Vonaesch Lab, the NCCR Microbiome consortium and Sara Mitri for fruitful discussions and valuable feedback on the study and manuscript. ChatGPT (OpenAI) was used to improve English and flow in the final text. Figures were created in part with BioRender. We further thank the Electron Microscopy Facility at the University of Lausanne for the help to image the phage particles.

## SUPPLEMENTARY DATA

**Supplementary Figure 1.**
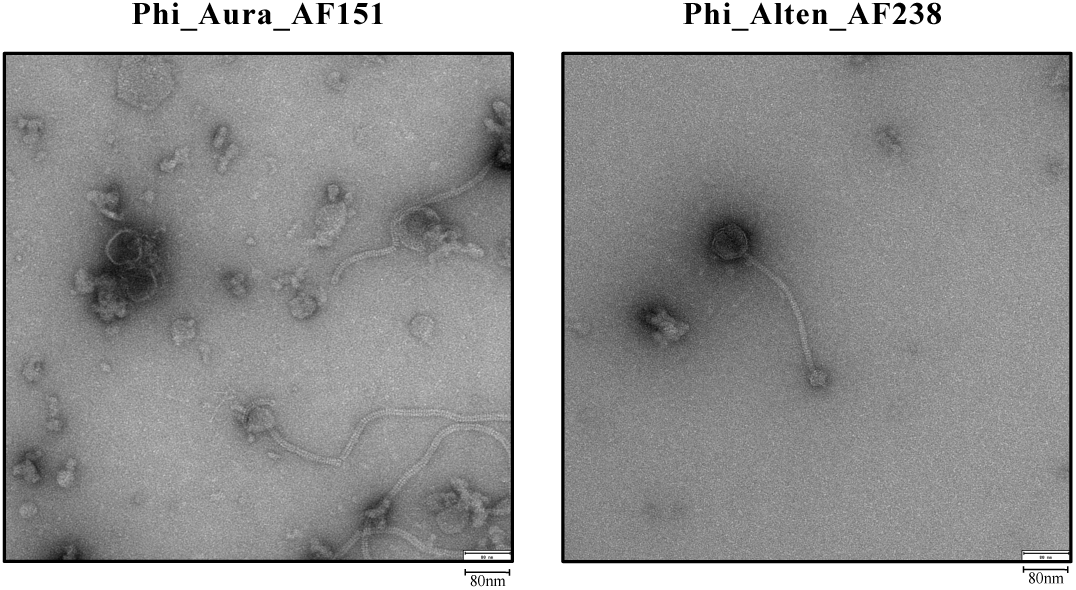
Transmission electron microscopy (TEM) images of phages phi_Aura_AF151 and phi_Alten_AF238. Phages were induced from *S. salivarius* strains AF151 and AF238, respectively. Both phages display *Siphovirus* morphology, characterized by long, non-contractile tails.

**Supplementary Figure 2.**
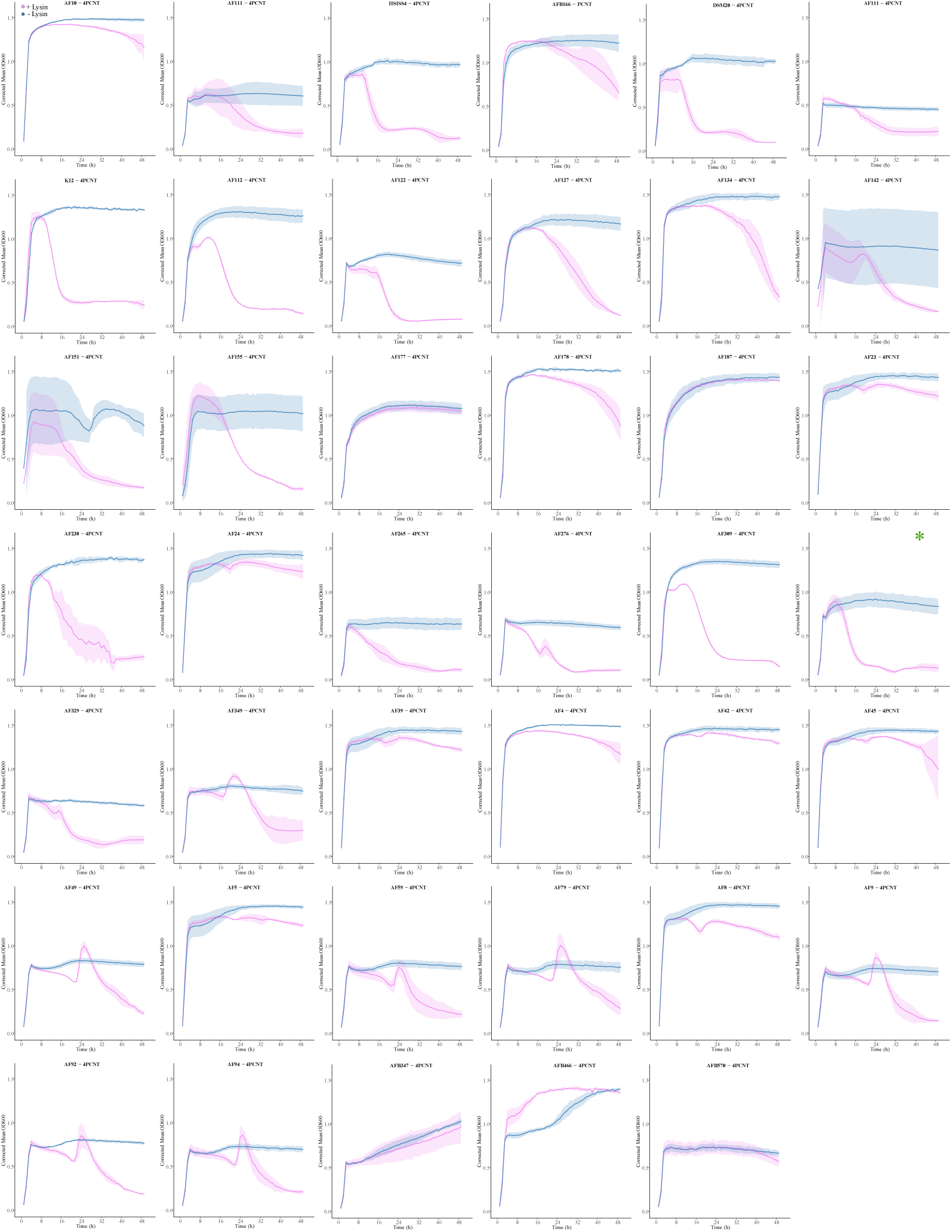
Growth inhibition of *S. salivarius* isolates by Lys_AF151 lysin. OD600nm growth curves over 48 hours for *S. salivarius* isolates grown in BHI medium with (Pink) or without (Blue) endolysin Lys_AF151 treatment. Each panel represents one isolate (strain name indicated above each plot). Growth was monitored every hour under anaerobic conditions. All experiments were performed in triplicates.

**Supplementary Figure 3.**
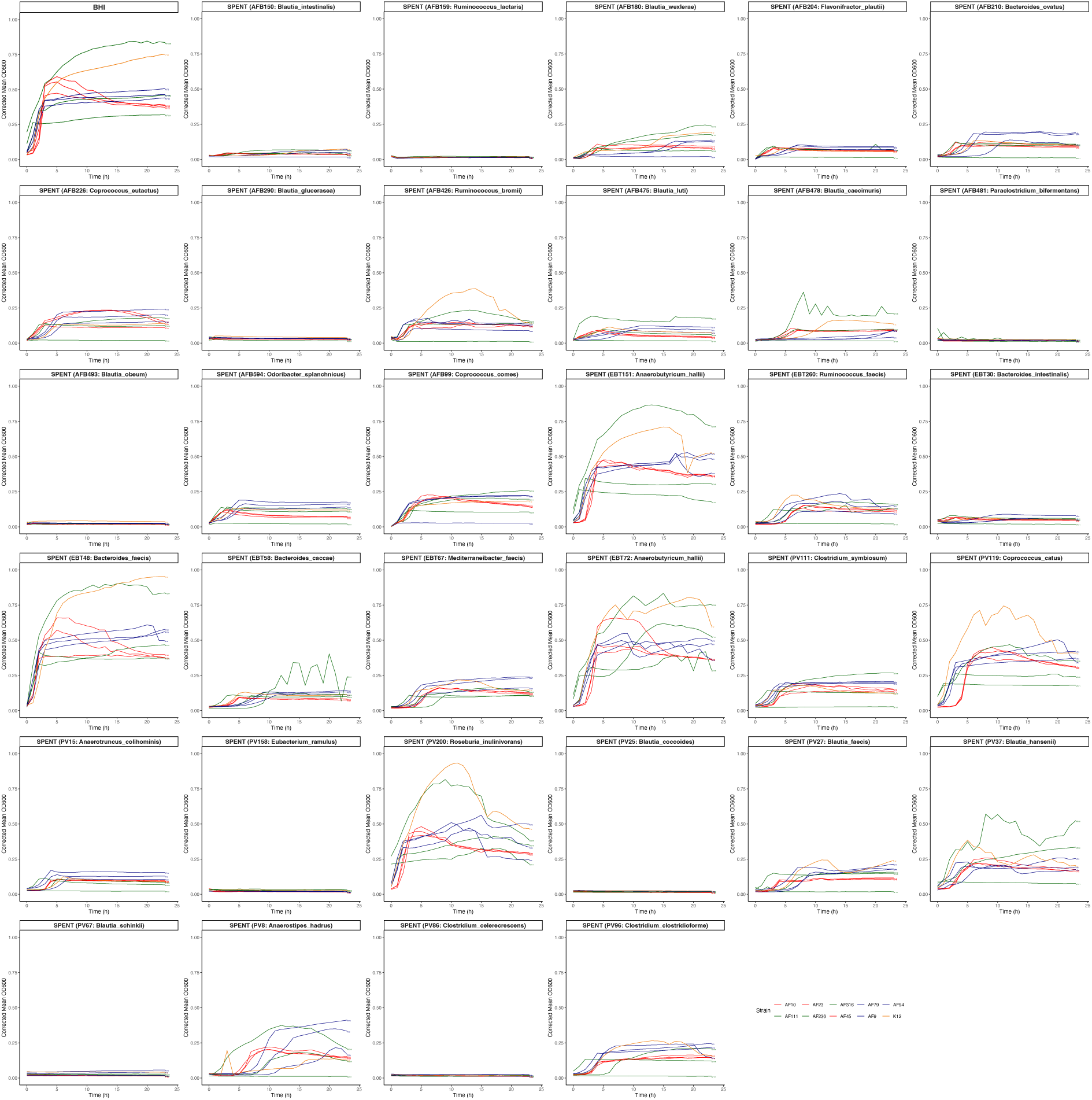
Results of the spent medium assay. The 10 selected *S. salivarius* strains were grown in fresh BHI for reference growth curves and grown in the spent medium from 33 “donor strains”. Each panel represents the growth in the spent medium of one specific donor strain. All experiments were performed twice with a single, representative experiment represented here. Colors for growth curves were assigned to *S. salivarius* strains according to the ANI cluster they belong (Red : cluster 1 = C1, Green : cluster 2 = C2, Blue : cluster 3 = C3, Orange : cluster 4 = C4

**Supplementary Table 1.**
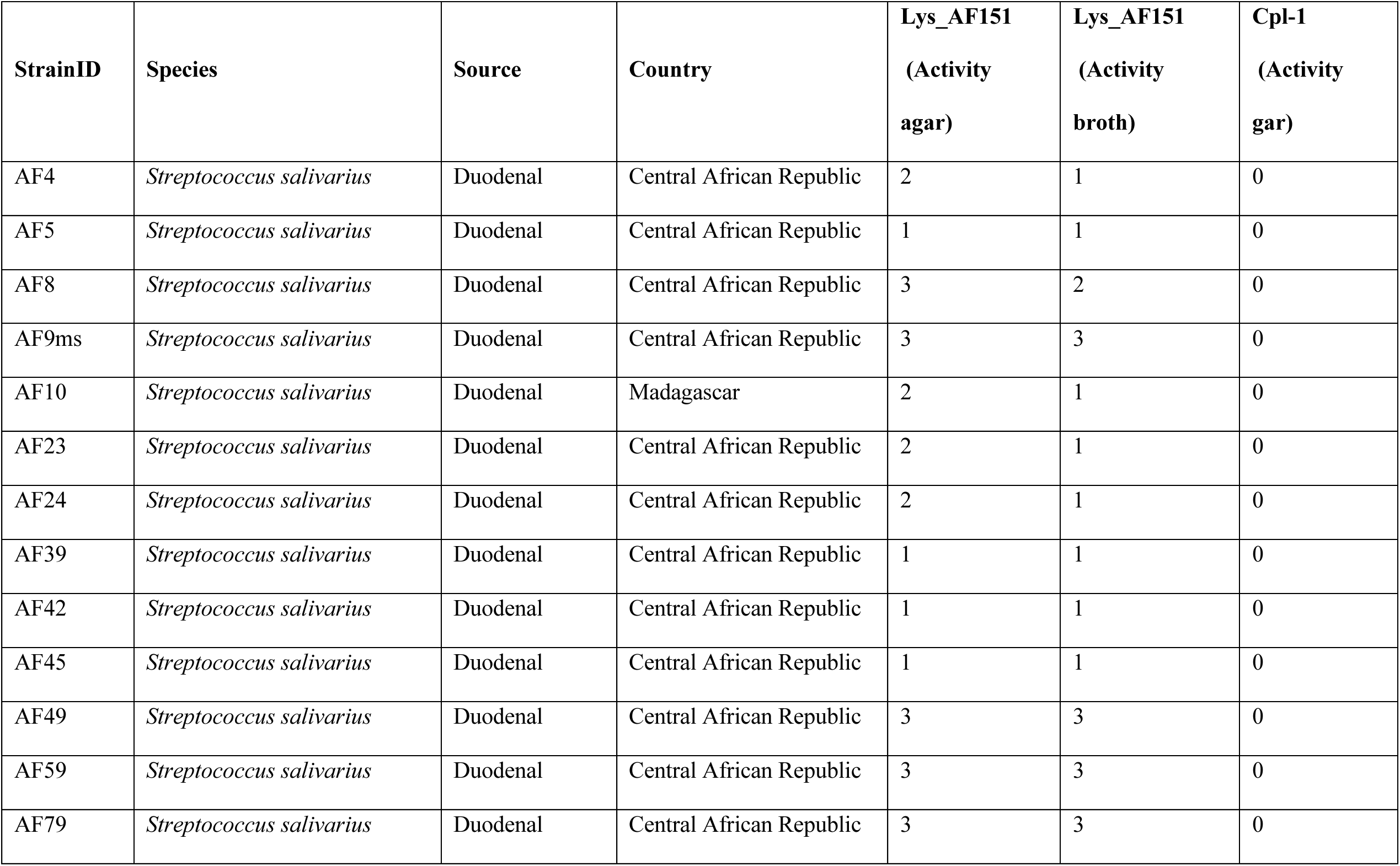
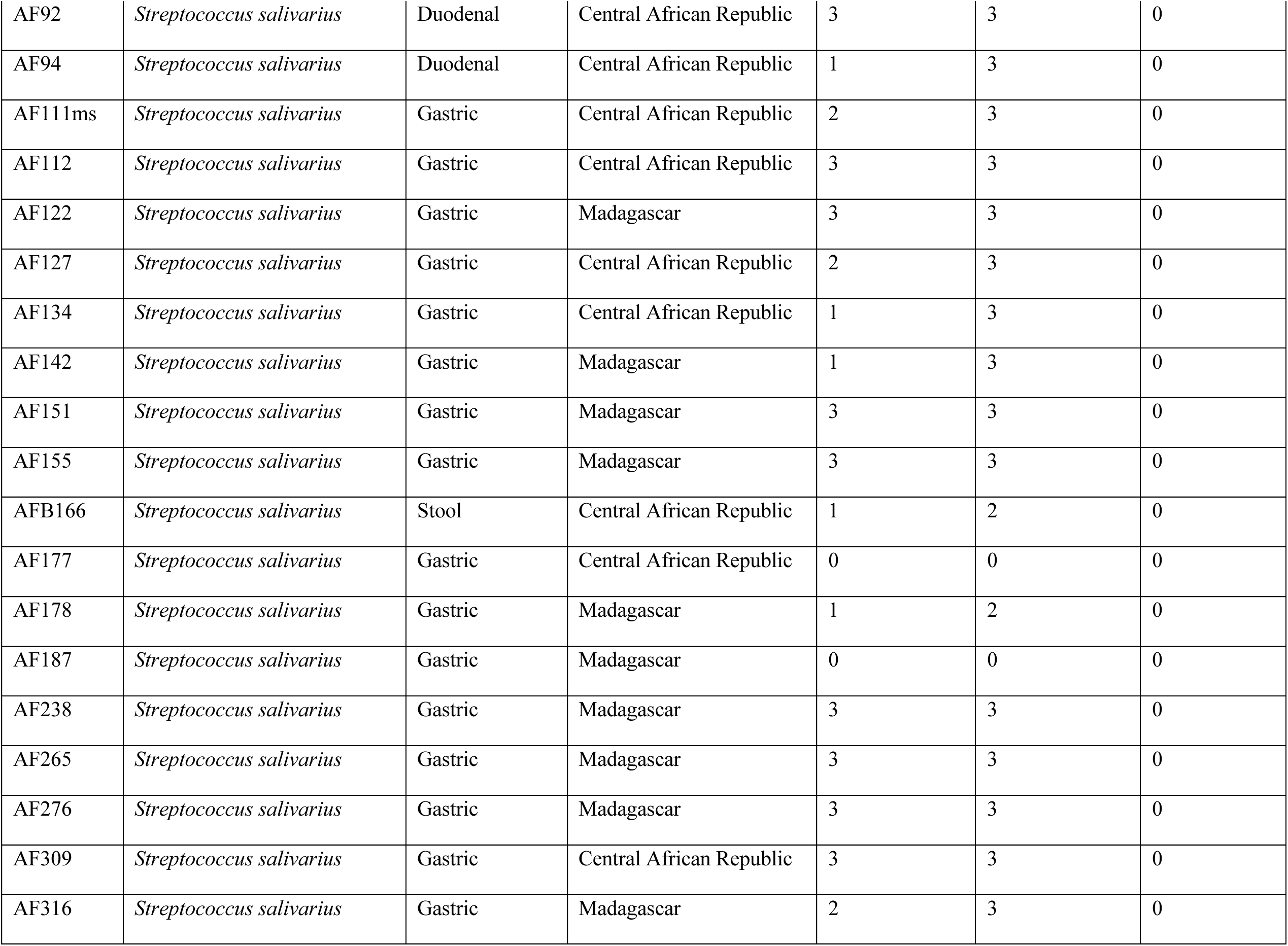
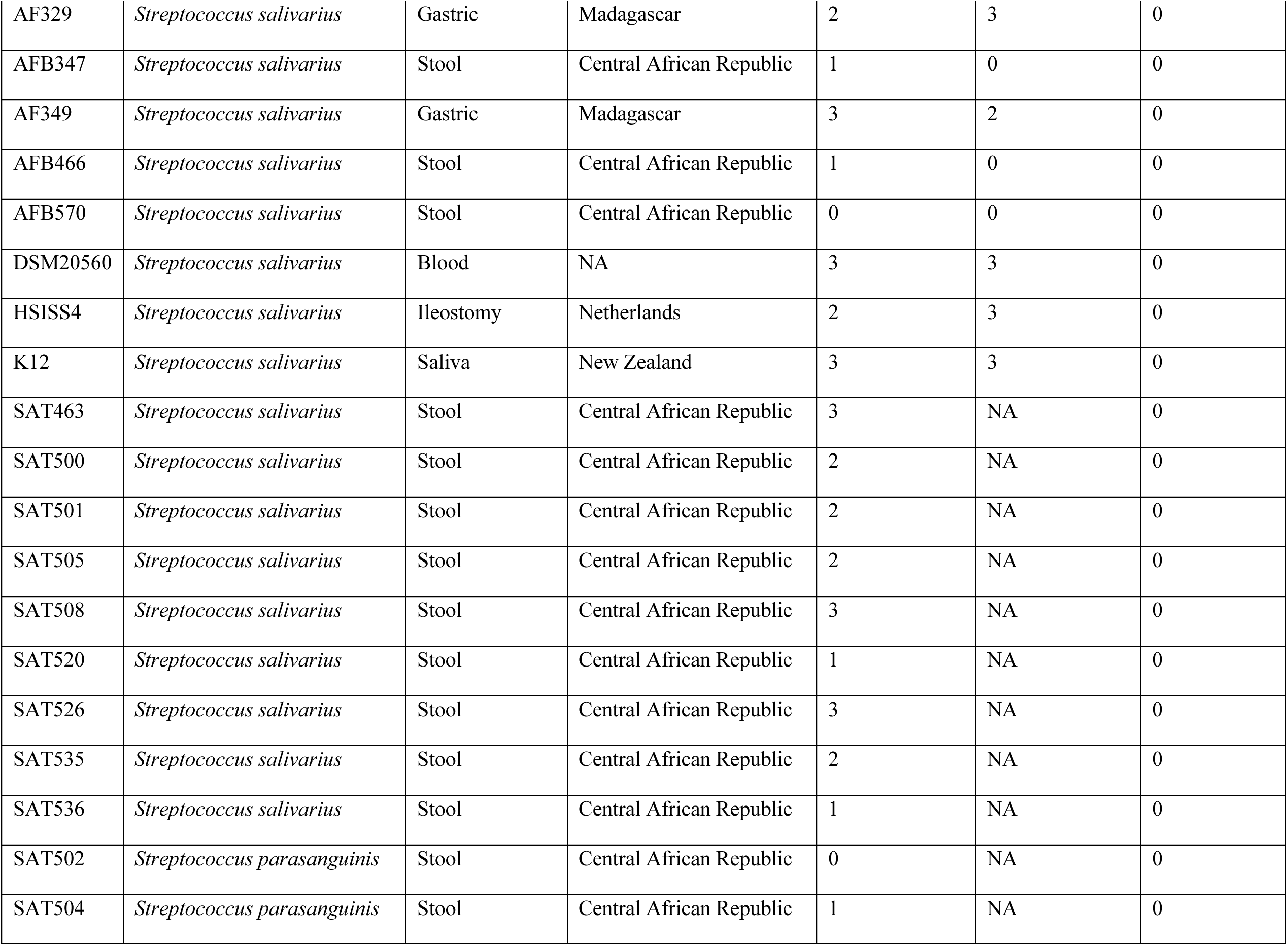
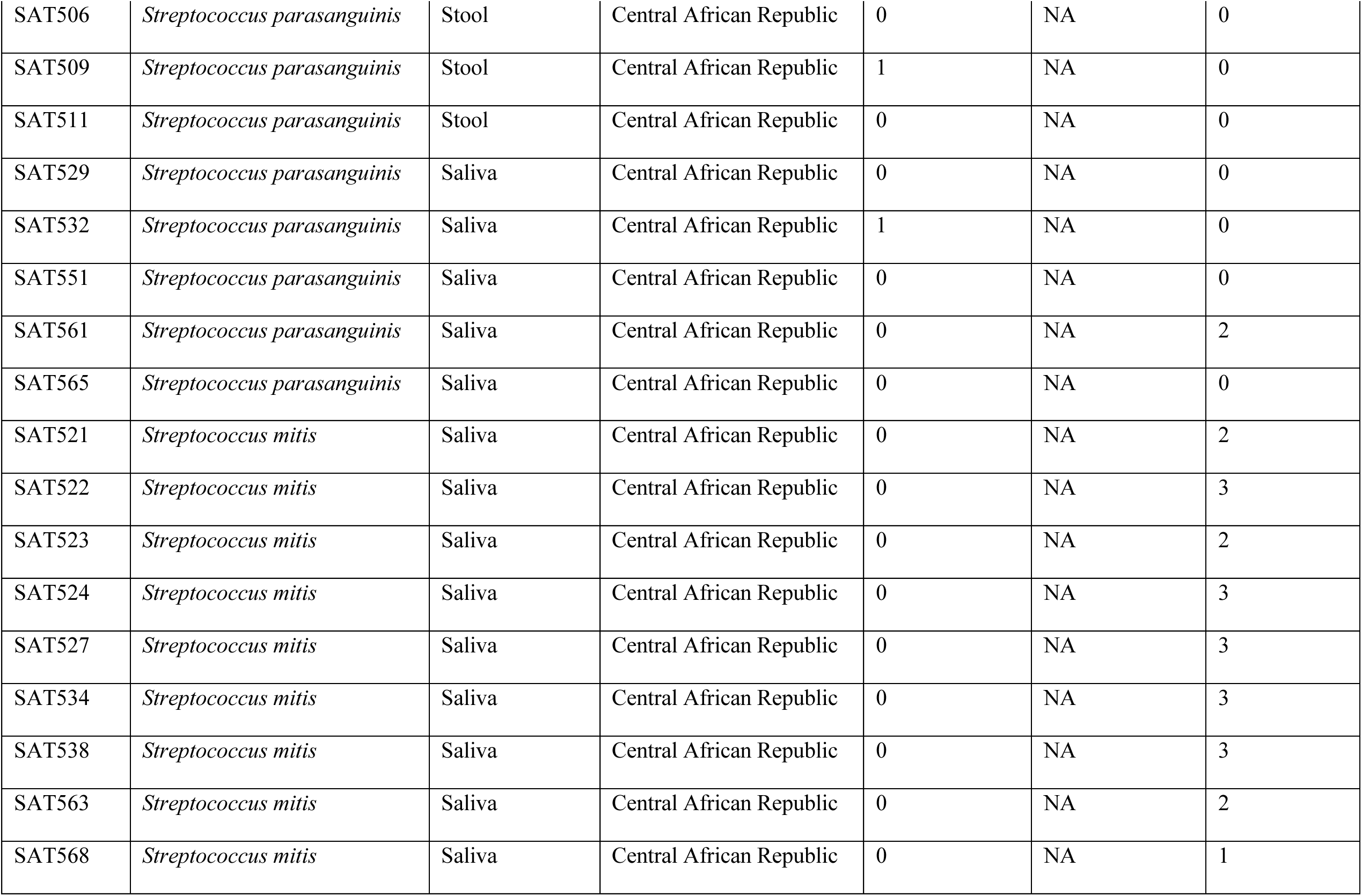
Strain used for the agar and broth lysin activity assays.

**Supplementary Table 2.**
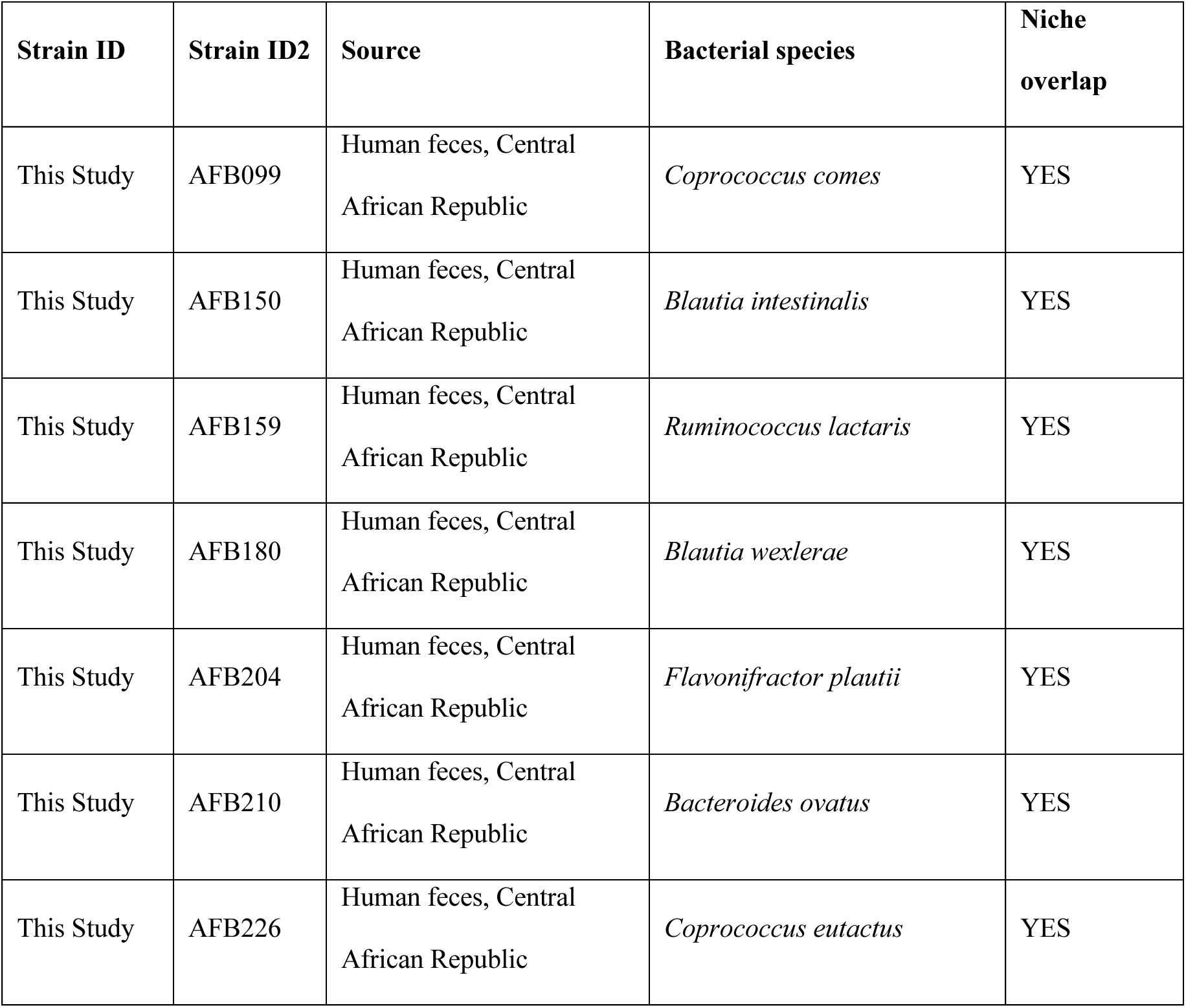
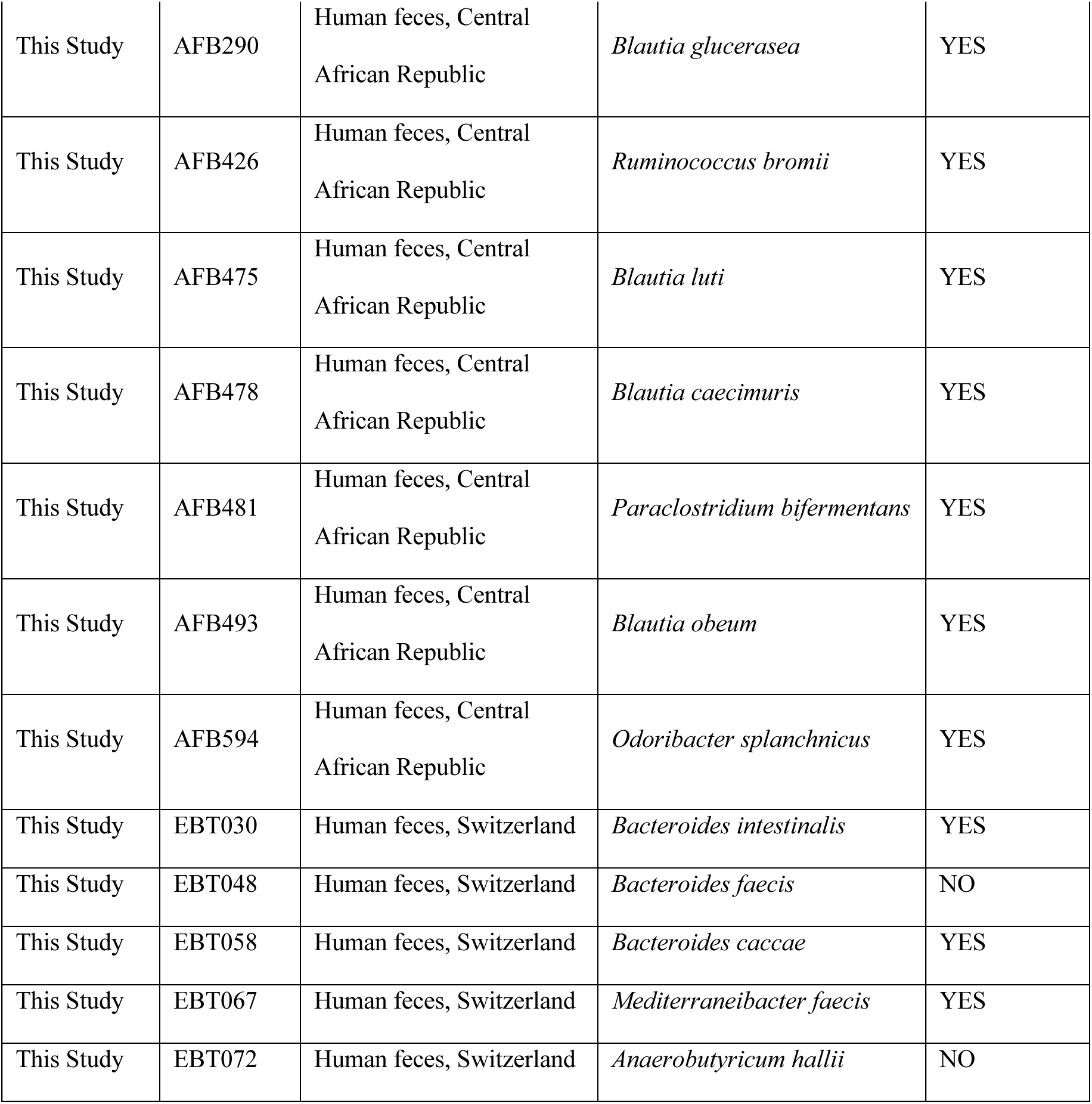
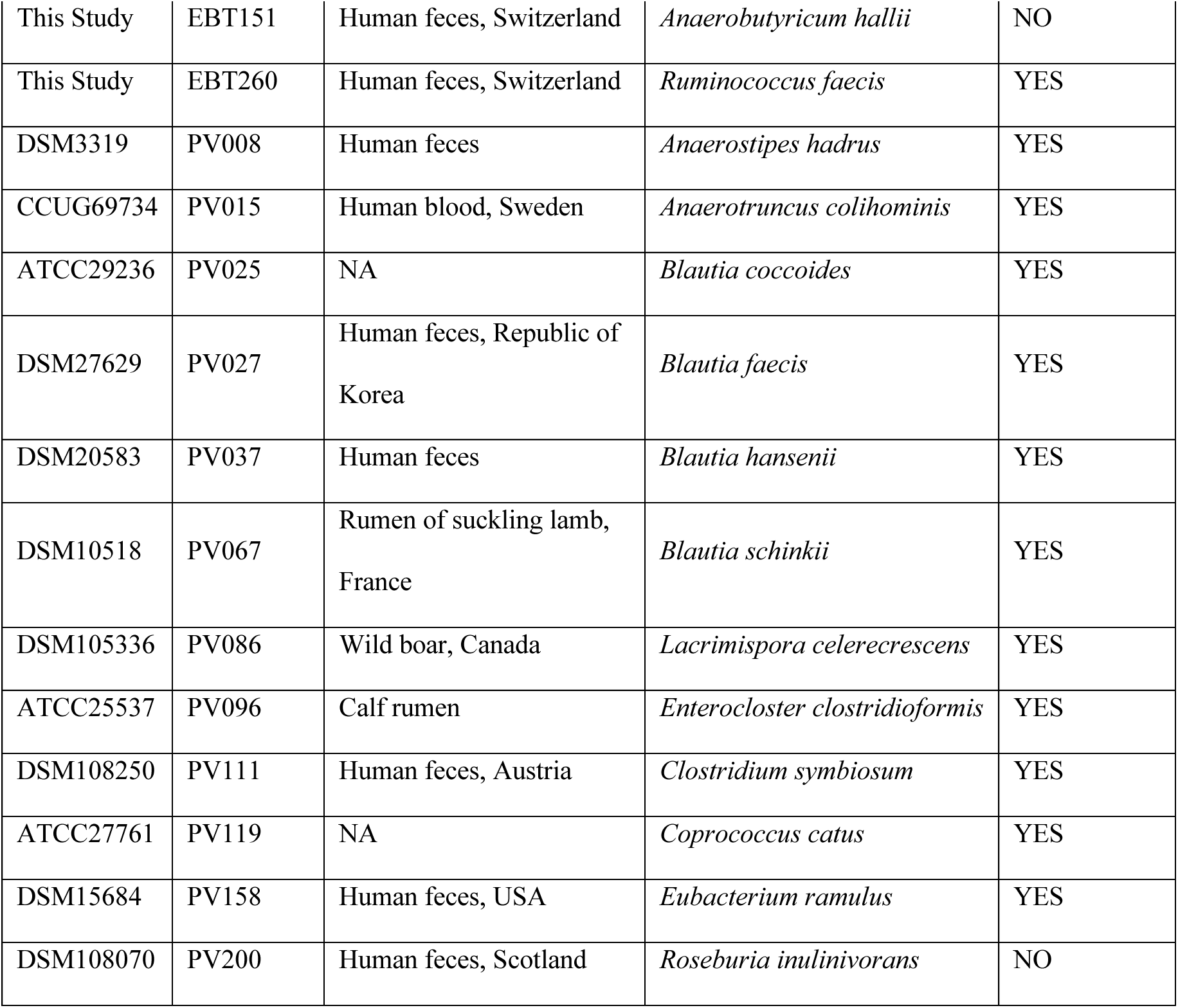
Strain used to generate the spent medium (“donor” strains)

**Supplementary Tables 3.**
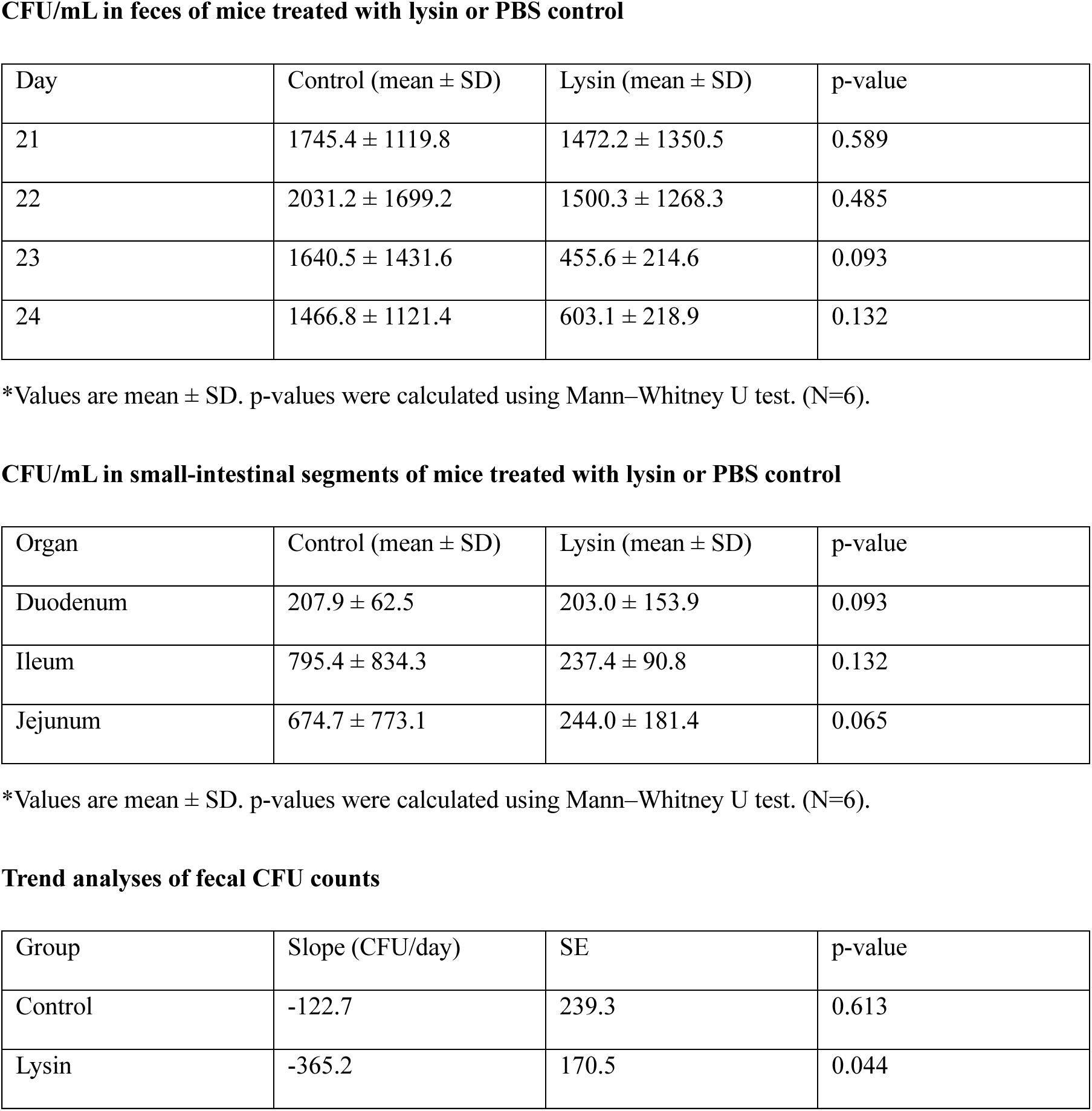
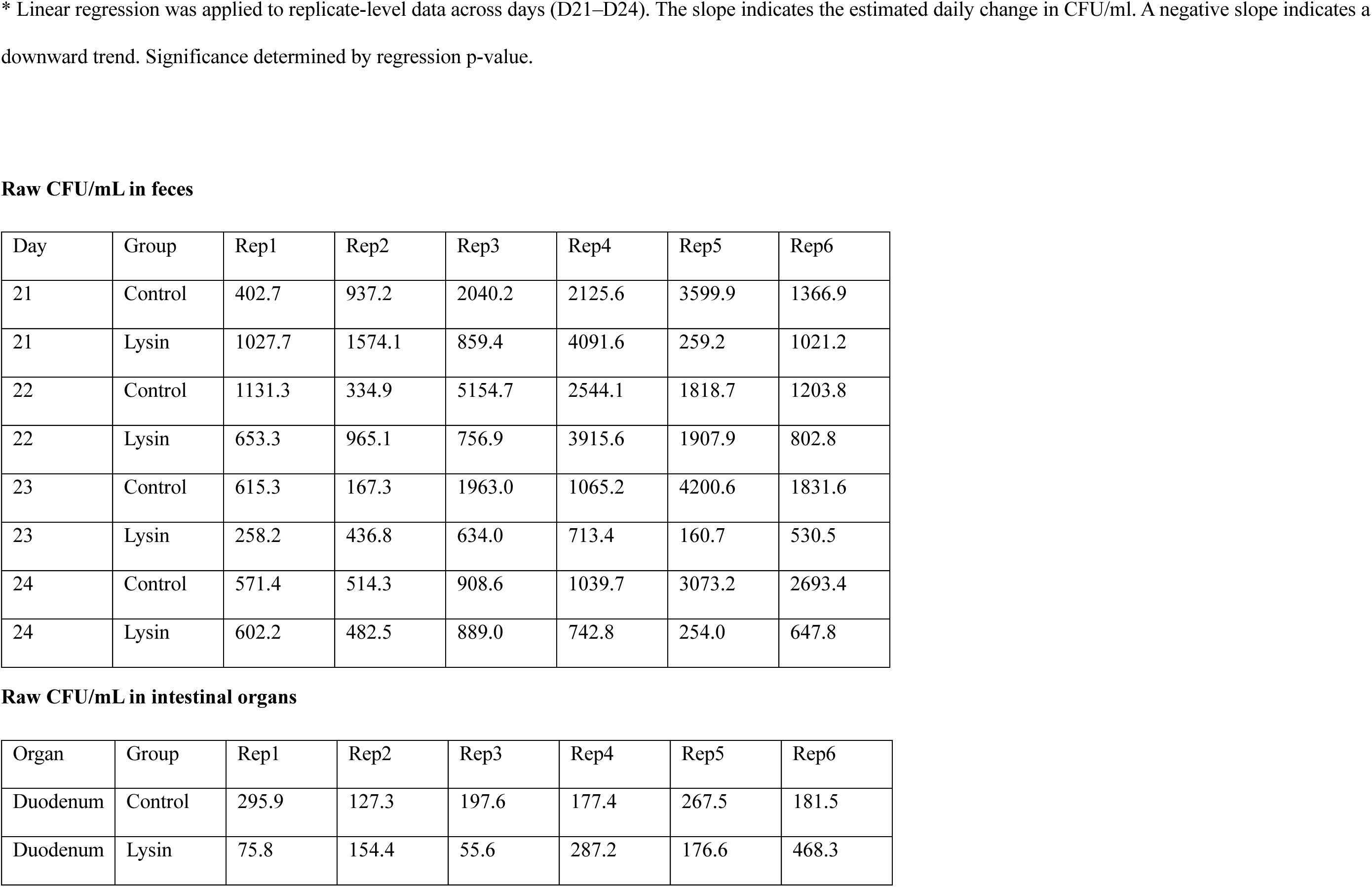
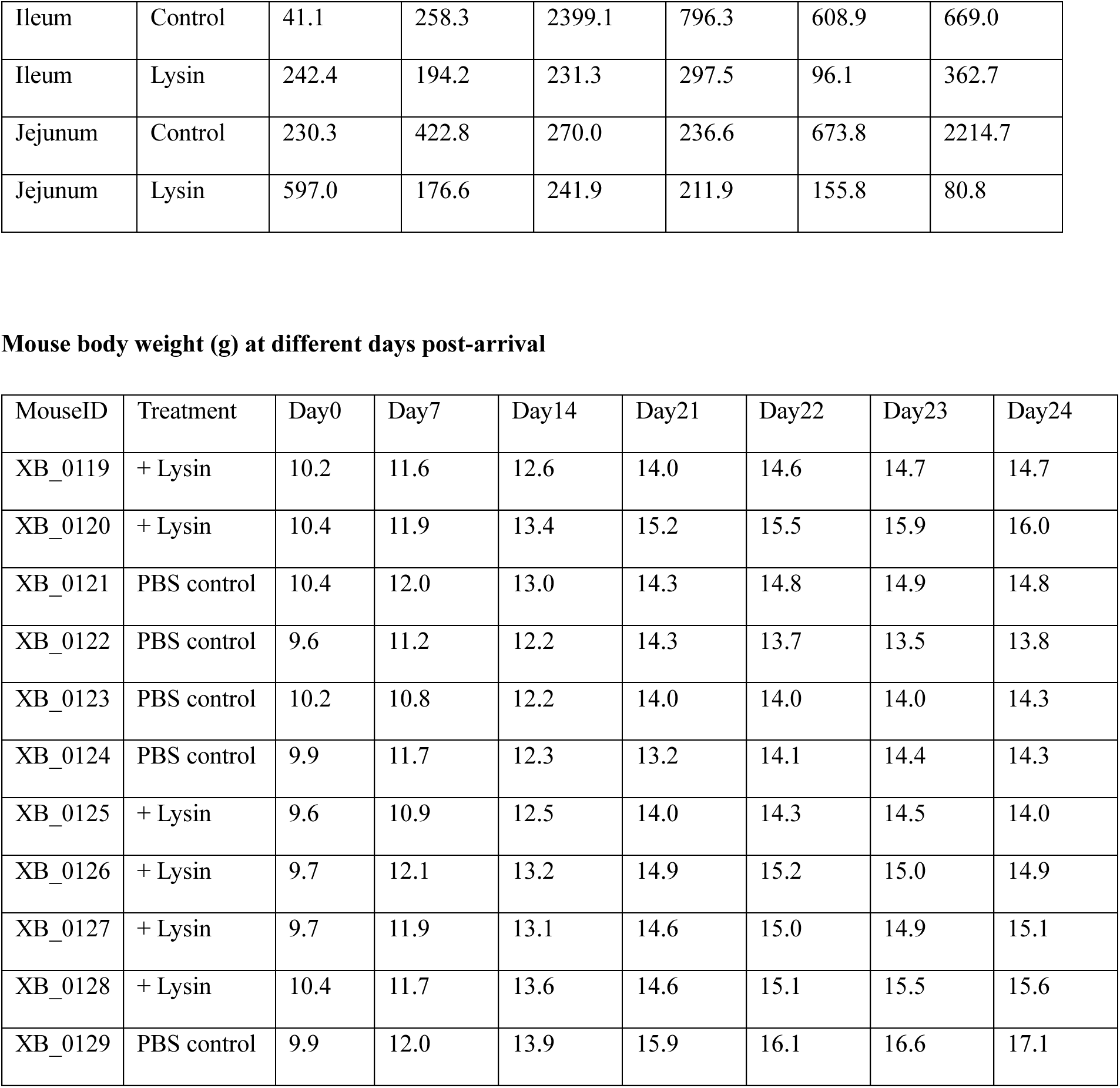
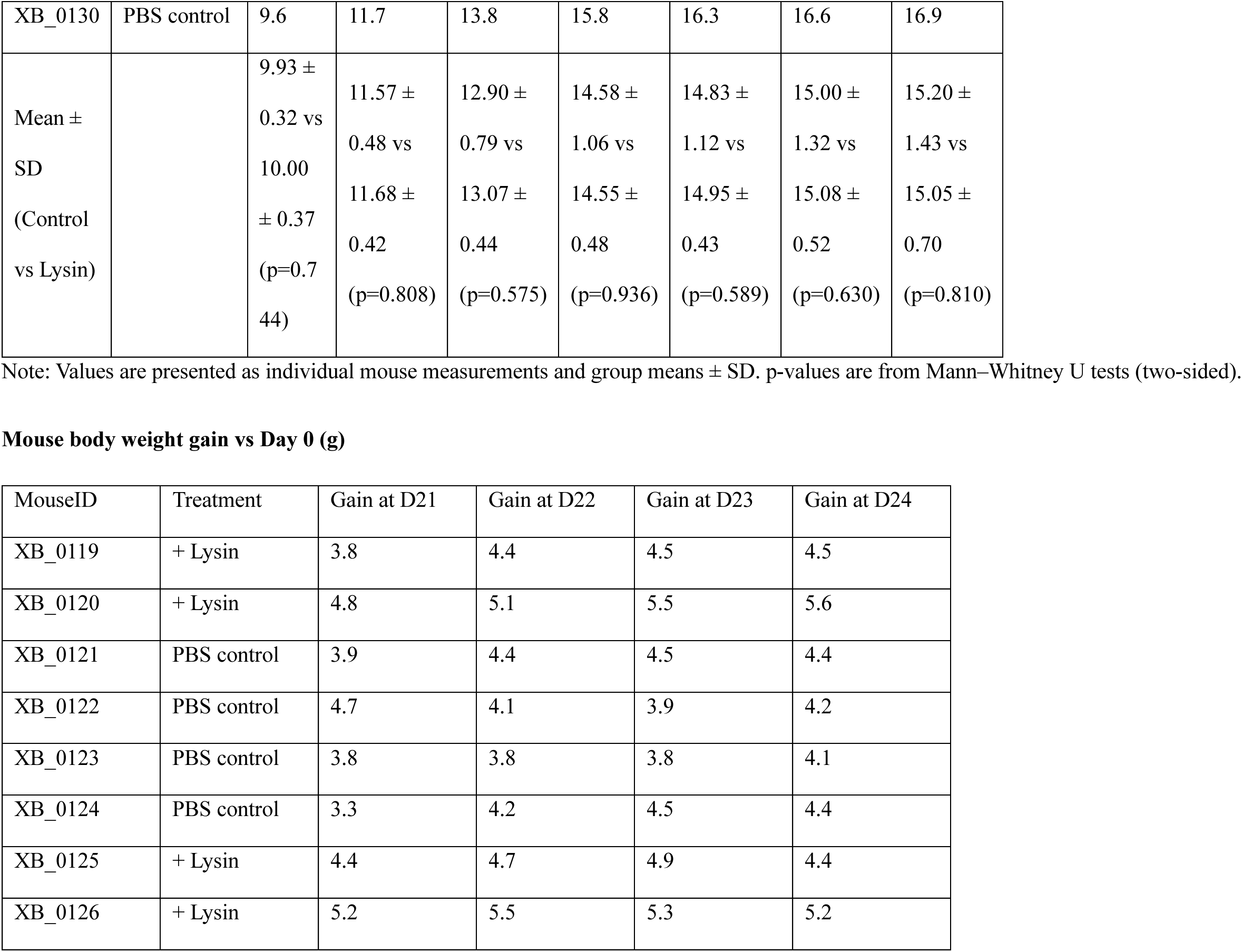
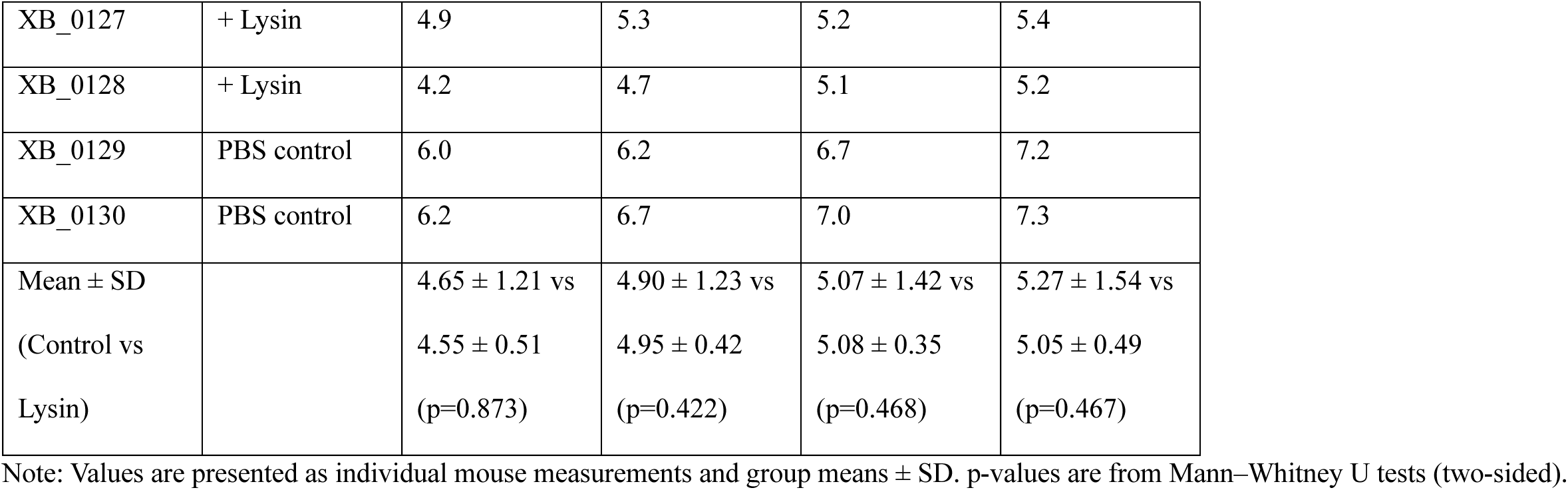
Mice related measurement and statistics.

## REFERENCES

1. Yersin, S. & Vonaesch, P. Small intestinal microbiota: from taxonomic composition to metabolism. Trends Microbiol. 32, 970–983 (2024).

2. Hernández-Cabanyero, C. & Vonaesch, P. Ectopic colonization by oral bacteria as an emerging theme in health and disease. FEMS Microbiol. Rev. 48, fuae012 (2024).

3. Vonaesch, P. et al. Stunted children display ectopic small intestinal colonization by oral bacteria, which cause lipid malabsorption in experimental models. Proc. Natl. Acad. Sci. U. S. A. 119, e2209589119 (2022).

4. Kastl, A. J., Terry, N. A., Wu, G. D. & Albenberg, L. G. The Structure and Function of the Human Small Intestinal Microbiota: Current Understanding and Future Directions. Cell. Mol. Gastroenterol. Hepatol. 9, 33–45 (2020).

5. El Aidy, S., van den Bogert, B. & Kleerebezem, M. The small intestine microbiota, nutritional modulation and relevance for health. Curr. Opin. Biotechnol. 32, 14–20 (2015).

6. Hou, K. et al. Microbiota in health and diseases. Signal Transduct. Target. Ther. 7, 135 (2022).

7. Chen, R. Y. et al. Duodenal Microbiota in Stunted Undernourished Children with Enteropathy. N. Engl. J. Med. 383, 321–333 (2020).

8. Prendergast, A. J. & Humphrey, J. H. The stunting syndrome in developing countries. Paediatr. Int. Child Health 34, 250–265 (2014).

9. Wells, J. C. et al. The double burden of malnutrition: aetiological pathways and consequences for health. Lancet Lond. Engl. 395, 75–88 (2020).

10. Subramanian, S. et al. Persistent gut microbiota immaturity in malnourished Bangladeshi children. Nature 510, 417–421 (2014).

11. Blanton, L. V. et al. Gut bacteria that prevent growth impairments transmitted by microbiota from malnourished children. Science 351, 10.1126/science.aad3311 aad3311 (2016).

12. Chibuye, M. et al. Systematic review of associations between gut microbiome composition and stunting in under-five children. NPJ Biofilms Microbiomes 10, 46 (2024).

13. Vonaesch, P. et al. Stunted childhood growth is associated with decompartmentalization of the gastrointestinal tract and overgrowth of oropharyngeal taxa. Proc. Natl. Acad. Sci. U. S. A. 115, E8489–E8498 (2018).

14. Collard, J.-M., et al. High prevalence of small intestine bacteria overgrowth and asymptomatic carriage of enteric pathogens in stunted children in Antananarivo, Madagascar. PLoS Negl. Trop. Dis. 16, e0009849 (2022).

15. Miao, P. et al. Exacerbation of allergic rhinitis by the commensal bacterium Streptococcus salivarius. Nat. Microbiol. 8, 218–230 (2023).

16. Kviatcovsky, D., Valdés-Mas, R., Federici, S. & Elinav, E. Phage therapy in noncommunicable diseases. Science 382, 266–267 (2023).

17. Federici, S., Nobs, S. P. & Elinav, E. Phages and their potential to modulate the microbiome and immunity. Cell. Mol. Immunol. 18, 889–904 (2021).

18. Duan, Y. et al. Bacteriophage targeting of gut bacterium attenuates alcoholic liver disease. Nature 575, 505–511 (2019).

19. Chou, W.-C., Huang, S.-C., Chiu, C.-H. & Chen, Y.-Y. M. YMC-2011, a Temperate Phage of Streptococcus salivarius 57.I. Appl. Environ. Microbiol. 83, e03186–16 (2017).

20. Suh, G. A. et al. Considerations for the Use of Phage Therapy in Clinical Practice. Antimicrob. Agents Chemother. 66, e0207121 (2022).

21. Heuler, J., Fortier, L.-C. & Sun, X. Clostridioides dihicile phage biology and application. FEMS Microbiol. Rev. 45, fuab012 (2021).

22. Umansky, A. A. & Fortier, L. C. The long and sinuous road to phage-based therapy of Clostridioides dihicile infections. Front. Med. 10, 1259427 (2023).

23. Pottie, I., Vázquez Fernández, R., Van de Wiele, T. & Briers, Y. Phage lysins for intestinal microbiome modulation: current challenges and enabling techniques. Gut Microbes 16, 2387144 (2024).

24. Fischetti, V. A. Bacteriophage lysins as ehective antibacterials. Curr. Opin. Microbiol. 11, 393–400 (2008).

25. Schuch, R. et al. Use of a bacteriophage lysin to identify a novel target for antimicrobial development. PloS One 8, e60754 (2013).

26. Oh, J., Warner, M., Ambler, J. E. & Schuch, R. The Lysin Exebacase Has a Low Propensity for Resistance Development in Staphylococcus aureus and Suppresses the Emergence of Resistance to Antistaphylococcal Antibiotics. Microbiol. Spectr. 11, e0526122 (2023).

27. Theuretzbacher, U., Outterson, K., Engel, A. & Karlén, A. The global preclinical antibacterial pipeline. Nat. Rev. Microbiol. 18, 275–285 (2020).

28. Cooper, C. J., Khan Mirzaei, M. & Nilsson, A. S. Adapting Drug Approval Pathways for Bacteriophage-Based Therapeutics. Front. Microbiol. 7, 1209 (2016).

29. Kaspar, U. et al. The Novel Phage-Derived Antimicrobial Agent HY-133 Is Active against Livestock-Associated Methicillin-Resistant Staphylococcus aureus. Antimicrob. Agents Chemother. 62, e00385–18 (2018).

30. Knaack, D. et al. Bactericidal activity of bacteriophage endolysin HY-133 against Staphylococcus aureus in comparison to other antibiotics as determined by minimum bactericidal concentrations and time-kill analysis. Diagn. Microbiol. Infect. Dis. 93, 362–368 (2019).

31. Raz, A., Serrano, A., Hernandez, A., Euler, C. W. & Fischetti, V. A. Isolation of Phage Lysins That Ehectively Kill Pseudomonas aeruginosa in Mouse Models of Lung and Skin Infection. Antimicrob. Agents Chemother. 63, e00024–19 (2019).

32. Shah, S. et al. Beyond antibiotics: phage-encoded lysins against Gram-negative pathogens. Front. Microbiol. 14, 1170418 (2023).

33. Vander Elst, N., Farmen, K., Knörr, L., Merlijn, L. & Iovino, F. Bacteriophage-derived endolysins restore antibiotic susceptibility in β-lactam- and macrolide-resistant Streptococcus pneumoniae infections. Mol. Med. Camb. Mass 31, 170 (2025).

34. Resch, G., Moreillon, P. & Fischetti, V. A. A stable phage lysin (Cpl-1) dimer with increased antipneumococcal activity and decreased plasma clearance. Int. J. Antimicrob. Agents 38, 516–521 (2011).

35. Oechslin, F., Daraspe, J., Giddey, M., Moreillon, P. & Resch, G. In vitro characterization of PlySK1249, a novel phage lysin, and assessment of its antibacterial activity in a mouse model of Streptococcus agalactiae bacteremia. Antimicrob. Agents Chemother. 57, 6276–6283 (2013).

36. Fowler, V. G. et al. Exebacase for patients with Staphylococcus aureus bloodstream infection and endocarditis. J. Clin. Invest. 130, 3750–3760 (2020).

37. Fowler, V. G. et al. Exebacase in Addition to Standard-of-Care Antibiotics for Staphylococcus aureus Bloodstream Infections and Right-Sided Infective Endocarditis: A Phase 3, Superiority-Design, Placebo-Controlled, Randomized Clinical Trial (DISRUPT). Clin. Infect. Dis. OJ. Publ. Infect. Dis. Soc. Am. 78, 1473–1481 (2024).

38. Wishart, D. S. et al. PHASTEST: faster than PHASTER, better than PHAST. Nucleic Acids Res. 51, W443–W450 (2023).

39. Oliveira, L., Tavares, P. & Alonso, J. C. Headful DNA packaging: bacteriophage SPP1 as a model system. Virus Res. 173, 247–259 (2013).

40. Garneau, J. R. et al. High-throughput identification of viral termini and packaging mechanisms in virome datasets using PhageTermVirome. Sci. Rep. 11, 18319 (2021).

41. Jones, P. et al. InterProScan 5: genome-scale protein function classification. Bioinforma. Oxf. Engl. 30, 1236–1240 (2014).

42. Hidalgo-Villeda, F. et al. Prolonged dysbiosis and altered immunity under nutritional intervention in a physiological mouse model of severe acute malnutrition. iScience 26, 106910 (2023).

43. Dos Santos, A. R., Di Martino, R., Testa, S. E. A. & Mitri, S. Classifying Interactions in a Synthetic Bacterial Community Is Hindered by Inhibitory Growth Medium. mSystems 7, e0023922 (2022).

44. Raman, A. S. et al. A sparse covarying unit that describes healthy and impaired human gut microbiota development. Science 365, eaau4735 (2019).

45. Ji, H. et al. Function analysis of choline binding domains (CBDs) of LytA, LytC and CbpD in biofilm formation of Streptococcus pneumoniae. Microb. Pathog. 174, 105939 (2023).

46. Torrens, G. & Cava, F. Mechanisms conferring bacterial cell wall variability and adaptivity. Biochem. Soc. Trans. 52, 1981–1993 (2024).

47. Yang, H. et al. Novel chimeric lysin with high-level antimicrobial activity against methicillin-resistant Staphylococcus aureus in vitro and in vivo. Antimicrob. Agents Chemother. 58, 536–542 (2014).

48. Murray, E., Draper, L. A., Ross, R. P. & Hill, C. The Advantages and Challenges of Using Endolysins in a Clinical Setting. Viruses 13, 680 (2021).

49. Aranda-Díaz, A. et al. Establishment and characterization of stable, diverse, fecal-derived in vitro microbial communities that model the intestinal microbiota. Cell Host Microbe 30, 260–272.e5 (2022).

50. Smith, D. R., Temime, L. & Opatowski, L. Microbiome-pathogen interactions drive epidemiological dynamics of antibiotic resistance: A modeling study applied to nosocomial pathogen control. eLife 10, e68764 (2021).

51. Colautti, A., Orecchia, E., Comi, G. & Iacumin, L. Lactobacilli, a Weapon to Counteract Pathogens through the Inhibition of Their Virulence Factors. J. Bacteriol. 204, e0027222 (2022).

52. Everard, A. et al. Cross-talk between Akkermansia muciniphila and intestinal epithelium controls diet-induced obesity. Proc. Natl. Acad. Sci. U. S. A. 110, 9066–9071 (2013).

53. Depommier, C. et al. Supplementation with Akkermansia muciniphila in overweight and obese human volunteers: a proof-of-concept exploratory study. Nat. Med. 25, 1096–1103 (2019).

54. Macchione, I. G. et al. Akkermansia muciniphila: key player in metabolic and gastrointestinal disorders. Eur. Rev. Med. Pharmacol. Sci. 23, 8075–8083 (2019).

55. VAN GYLSWYK, N. O. & VAN DER TOORN, J. J. T. K. Eubacterium uniforme sp. nov. and Eubacterium xylanophilum sp. nov., Fiber-Digesting Bacteria from the Rumina of Sheep Fed Corn Stover. Int. J. Syst. Evol. Microbiol. 35, 323–326 (1985).

56. Ze, X., Duncan, S. H., Louis, P. & Flint, H. J. Ruminococcus bromii is a keystone species for the degradation of resistant starch in the human colon. ISME J. 6, 1535–1543 (2012).

57. Pascal, V. et al. A microbial signature for Crohn’s disease. Gut 66, 813–822 (2017).

58. Bauer, E., Laczny, C. C., Magnusdottir, S., Wilmes, P. & Thiele, I. Phenotypic diherentiation of gastrointestinal microbes is reflected in their encoded metabolic repertoires. Microbiome 3, 55 (2015).

59. Schmidt, T. S. B. et al. Drivers and determinants of strain dynamics following fecal microbiota transplantation. Nat. Med. 28, 1902–1912 (2022).

60. Goyal, A., Bittleston, L. S., Leventhal, G. E., Lu, L. & Cordero, O. X. Interactions between strains govern the eco-evolutionary dynamics of microbial communities. eLife 11, e74987 (2022).

61. Harhala, M. et al. Safety Studies of Pneumococcal Endolysins Cpl-1 and Pal. Viruses 10, 638 (2018).

62. Cheng, M. et al. Endolysin LysEF-P10 shows potential as an alternative treatment strategy for multidrug-resistant Enterococcus faecalis infections. Sci. Rep. 7, 10164 (2017).

63. Barber, J. N. et al. Species interactions constrain adaptation and preserve ecological stability in an experimental microbial community. ISME J. 16, 1442–1452 (2022).

64. Gerstmans, H. et al. A VersaTile-driven platform for rapid hit-to-lead development of engineered lysins. Sci. Adv. 6, eaaz1136 (2020).

65. Gerstmans, H. et al. Distinct mode of action of a highly stable, engineered phage lysin killing Gram-negative bacteria. Microbiol. Spectr. 11, e0181323 (2023).

66. Vonaesch, P. et al. Identifying the etiology and pathophysiology underlying stunting and environmental enteropathy: study protocol of the AFRIBIOTA project. BMC Pediatr. 18, 236 (2018).

67. Asare, P. T. et al. A MALDI-TOF MS library for rapid identification of human commensal gut bacteria from the class Clostridia. Front. Microbiol. 14, 1104707 (2023).

68. Chen, S., Zhou, Y., Chen, Y. & Gu, J. fastp: an ultra-fast all-in-one FASTQ preprocessor. Bioinforma. Oxf. Engl. 34, i884–i890 (2018).

69. Bankevich, A. et al. SPAdes: a new genome assembly algorithm and its applications to single-cell sequencing. J. Comput. Biol. J. Comput. Mol. Cell Biol. 19, 455–477 (2012).

70. Seemann, T. Prokka: rapid prokaryotic genome annotation. Bioinforma. Oxf. Engl. 30, 2068–2069 (2014).

71. Langmead, B. & Salzberg, S. L. Fast gapped-read alignment with Bowtie 2. Nat. Methods 9, 357–359 (2012).

72. Schmidt, B. M. et al. Clinker: visualizing fusion genes detected in RNA-seq data. GigaScience 7, giy079 (2018).

73. Blum, M. et al. The InterPro protein families and domains database: 20 years on. Nucleic Acids Res. 49, D344–D354 (2021).

74. Callahan, B. J. et al. DADA2: High-resolution sample inference from Illumina amplicon data. Nat. Methods 13, 581–583 (2016).

75. Pruesse, E. et al. SILVA: a comprehensive online resource for quality checked and aligned ribosomal RNA sequence data compatible with ARB. Nucleic Acids Res. 35, 7188–7196 (2007).

76. McMurdie, P. J. & Holmes, S. Phyloseq: a bioconductor package for handling and analysis of high-throughput phylogenetic sequence data. Pac. Symp. Biocomput. Pac. Symp. Biocomput. 235–246 (2012).

77. Ghezzi, H., et al. PUPpy: a primer design pipeline for substrain-level microbial detection and absolute quantification. mSphere 9, e0036024 (2024).

