## Supplementary Information for "Phage Endolysin Enables Targeted Manipulation of the Small Intestinal Microbiota and Uncovers Niche Overlap Between Oral and Butyrate-Producing Taxa"

[+] Co-last authors

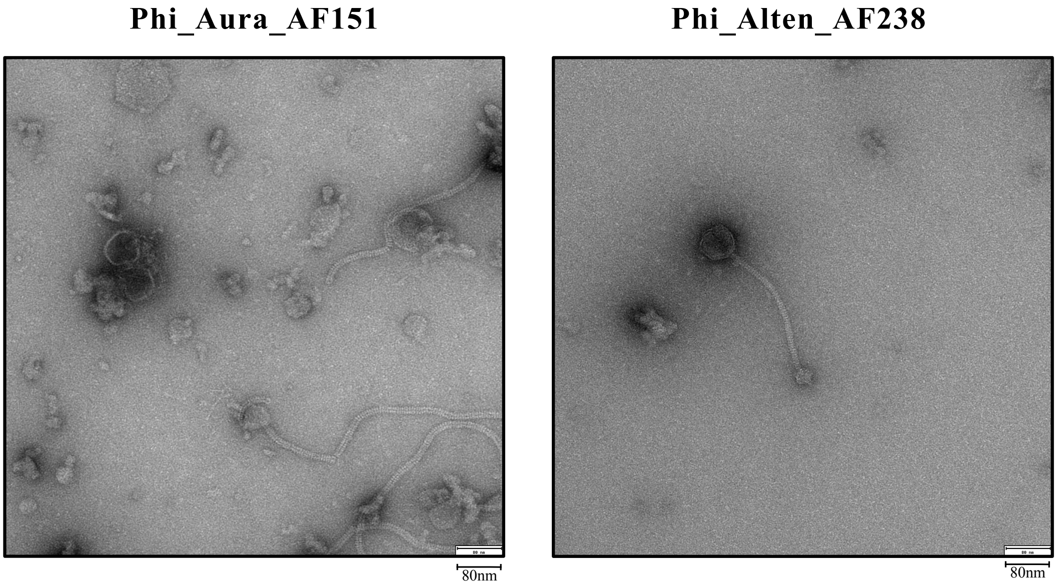


**Supplementary Figure 1. Transmission electron microscopy (TEM) images of phages phi_Aura_AF151 and phi_Alten_AF238.** Phages were induced from *S. salivarius* strains AF151 and AF238, respectively. Both phages display *Siphovirus* morphology, characterized by long, non-contractile tails.

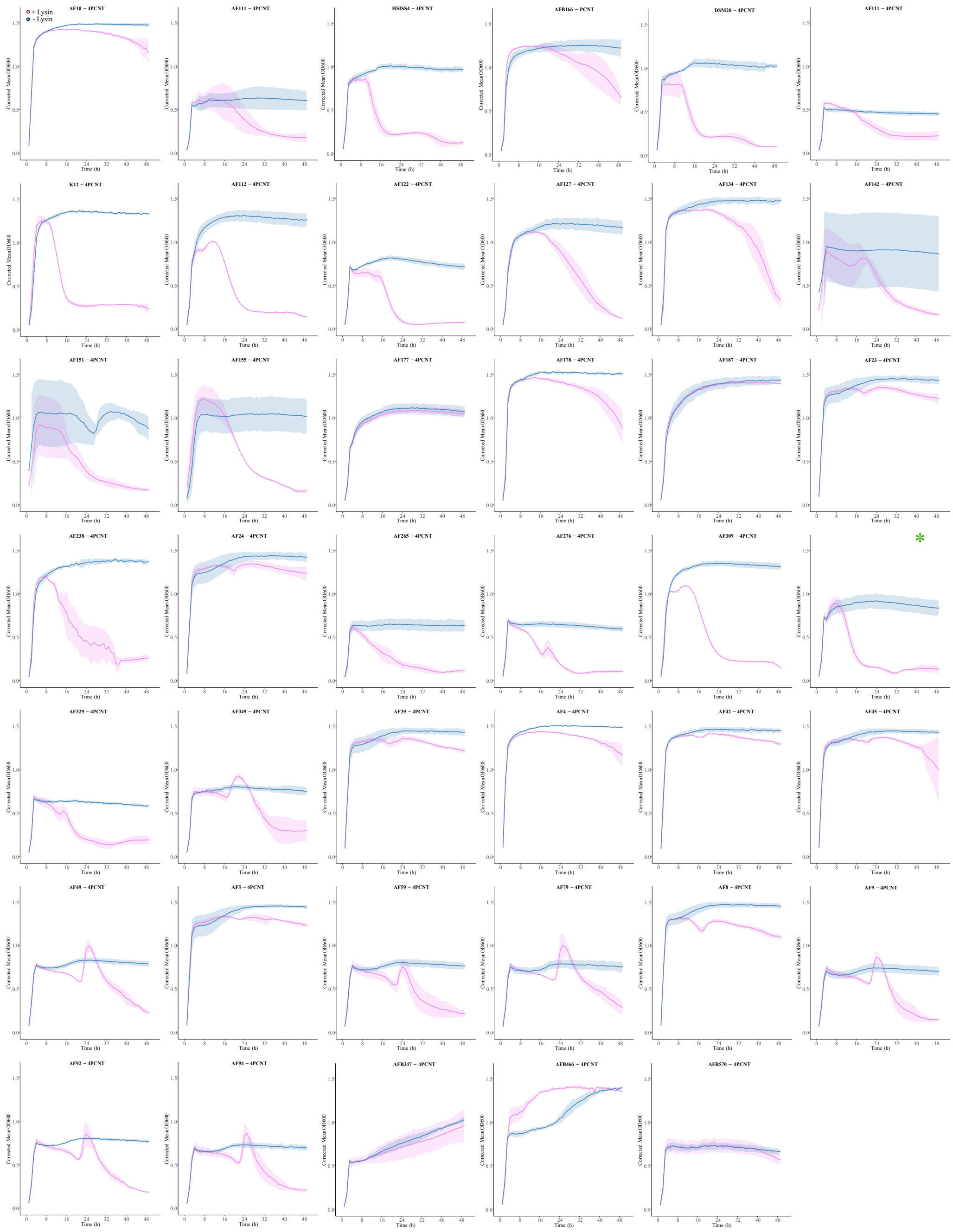


**Supplementary Figure 2. Growth inhibition of *S. salivarius* isolates by Lys_AF151 lysin.**

OD600nm growth curves over 48 hours for *S. salivarius* isolates grown in BHI medium with (Pink) or without (Blue) endolysin Lys_AF151 treatment. Each panel represents one isolate (strain name indicated above each plot). Growth was monitored every hour under anaerobic conditions. All experiments were performed in triplicates.

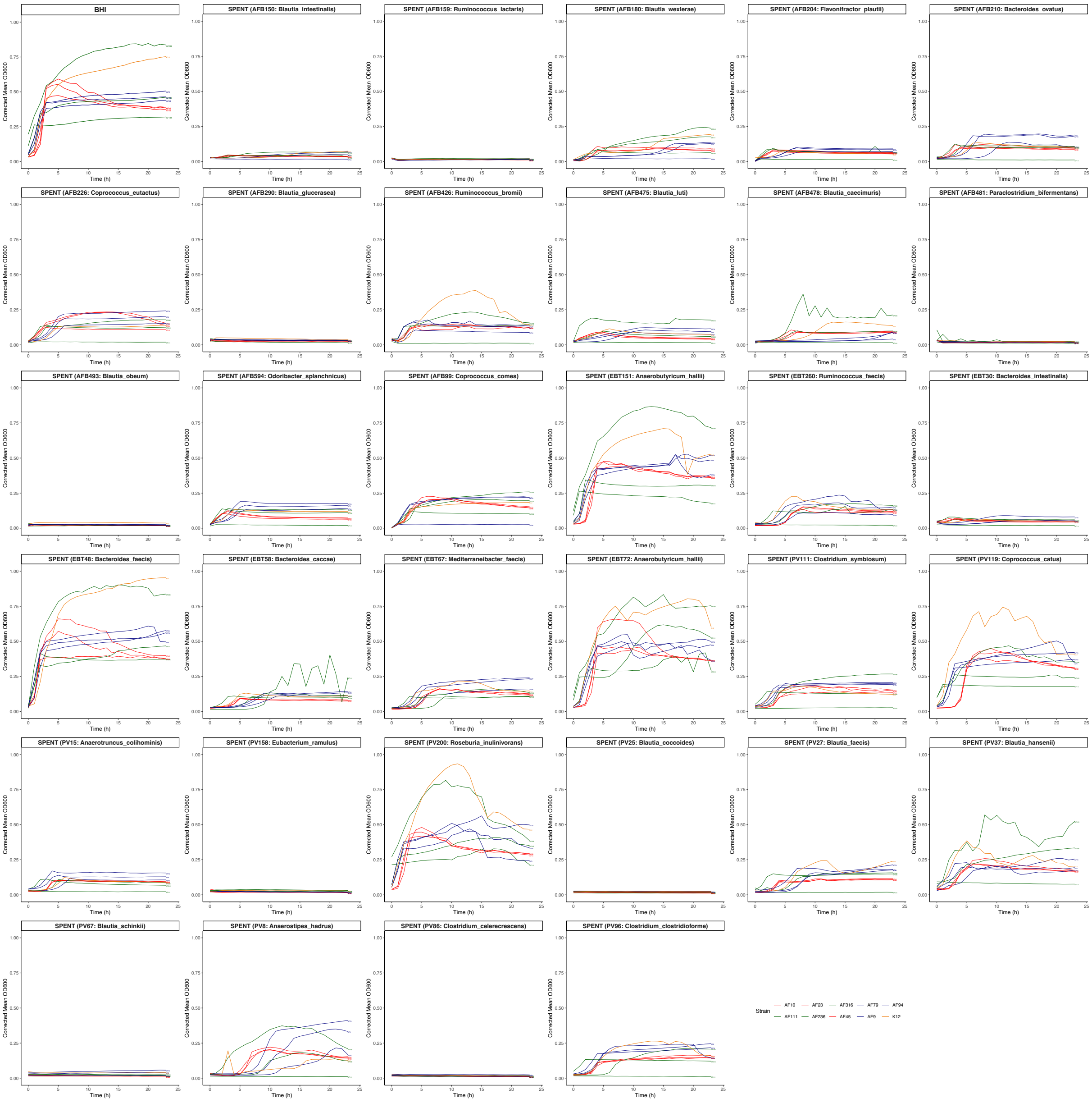


**Supplementary Figure 3.** Results of the spent medium assay. The 10 selected *S. salivarius* strains were grown in fresh BHI for reference growth curves and grown in the spent medium from 33 "donor strains". Each panel represents the growth in the spent medium of one specific donor strain. All experiments were performed twice with a single, representative experiment represented here. Colors for growth curves were assigned to *S. salivarius* strains according to the ANI cluster they belong (Red : cluster 1 = C1, Green : cluster 2 = C2, Blue : cluster 3 = C3, Orange : cluster 4 = C4

**Supplementary Table 1.** Strain used for the agar and broth lysin activity assays.

| **StrainID** | **Species** | **Source** | **Country** | **Lys_AF151 (Activity agar)** | **Lys_AF151 (Activity broth)** | **Cpl-1 (Activity gar)** |
|---|---|---|---|---|---|---|
| AF4 | *Streptococcus salivarius* | Duodenal | Central African Republic | 2 | 1 | 0 |
| AF5 | *Streptococcus salivarius* | Duodenal | Central African Republic | 1 | 1 | 0 |
| AF8 | *Streptococcus salivarius* | Duodenal | Central African Republic | 3 | 2 | 0 |
| AF9ms | *Streptococcus salivarius* | Duodenal | Central African Republic | 3 | 3 | 0 |
| AF10 | *Streptococcus salivarius* | Duodenal | Madagascar | 2 | 1 | 0 |
| AF23 | *Streptococcus salivarius* | Duodenal | Central African Republic | 2 | 1 | 0 |
| AF24 | *Streptococcus salivarius* | Duodenal | Central African Republic | 2 | 1 | 0 |
| AF39 | *Streptococcus salivarius* | Duodenal | Central African Republic | 1 | 1 | 0 |
| AF42 | *Streptococcus salivarius* | Duodenal | Central African Republic | 1 | 1 | 0 |
| AF45 | *Streptococcus salivarius* | Duodenal | Central African Republic | 1 | 1 | 0 |
| AF49 | *Streptococcus salivarius* | Duodenal | Central African Republic | 3 | 3 | 0 |
| AF59 | *Streptococcus salivarius* | Duodenal | Central African Republic | 3 | 3 | 0 |
| AF79 | *Streptococcus salivarius* | Duodenal | Central African Republic | 3 | 3 | 0 |

| | | | | | | |
|---|---|---|---|---|---|---|
| AF92 | *Streptococcus salivarius* | Duodenal | Central African Republic | 3 | 3 | 0 |
| AF94 | *Streptococcus salivarius* | Duodenal | Central African Republic | 1 | 3 | 0 |
| AF111ms | *Streptococcus salivarius* | Gastric | Central African Republic | 2 | 3 | 0 |
| AF112 | *Streptococcus salivarius* | Gastric | Central African Republic | 3 | 3 | 0 |
| AF122 | *Streptococcus salivarius* | Gastric | Madagascar | 3 | 3 | 0 |
| AF127 | *Streptococcus salivarius* | Gastric | Central African Republic | 2 | 3 | 0 |
| AF134 | *Streptococcus salivarius* | Gastric | Central African Republic | 1 | 3 | 0 |
| AF142 | *Streptococcus salivarius* | Gastric | Madagascar | 1 | 3 | 0 |
| AF151 | *Streptococcus salivarius* | Gastric | Madagascar | 3 | 3 | 0 |
| AF155 | *Streptococcus salivarius* | Gastric | Madagascar | 3 | 3 | 0 |
| AFB166 | *Streptococcus salivarius* | Stool | Central African Republic | 1 | 2 | 0 |
| AF177 | *Streptococcus salivarius* | Gastric | Central African Republic | 0 | 0 | 0 |
| AF178 | *Streptococcus salivarius* | Gastric | Madagascar | 1 | 2 | 0 |
| AF187 | *Streptococcus salivarius* | Gastric | Madagascar | 0 | 0 | 0 |
| AF238 | *Streptococcus salivarius* | Gastric | Madagascar | 3 | 3 | 0 |
| AF265 | *Streptococcus salivarius* | Gastric | Madagascar | 3 | 3 | 0 |
| AF276 | *Streptococcus salivarius* | Gastric | Madagascar | 3 | 3 | 0 |
| AF309 | *Streptococcus salivarius* | Gastric | Central African Republic | 3 | 3 | 0 |
| AF316 | *Streptococcus salivarius* | Gastric | Madagascar | 2 | 3 | 0 |

| | | | | | | |
|---|---|---|---|---|---|---|
| AF329 | *Streptococcus salivarius* | Gastric | Madagascar | 2 | 3 | 0 |
| AFB347 | *Streptococcus salivarius* | Stool | Central African Republic | 1 | 0 | 0 |
| AF349 | *Streptococcus salivarius* | Gastric | Madagascar | 3 | 2 | 0 |
| AFB466 | *Streptococcus salivarius* | Stool | Central African Republic | 1 | 0 | 0 |
| AFB570 | *Streptococcus salivarius* | Stool | Central African Republic | 0 | 0 | 0 |
| DSM20560 | *Streptococcus salivarius* | Blood | NA | 3 | 3 | 0 |
| HSISS4 | *Streptococcus salivarius* | Ileostomy | Netherlands | 2 | 3 | 0 |
| K12 | *Streptococcus salivarius* | Saliva | New Zealand | 3 | 3 | 0 |
| SAT463 | *Streptococcus salivarius* | Stool | Central African Republic | 3 | NA | 0 |
| SAT500 | *Streptococcus salivarius* | Stool | Central African Republic | 2 | NA | 0 |
| SAT501 | *Streptococcus salivarius* | Stool | Central African Republic | 2 | NA | 0 |
| SAT505 | *Streptococcus salivarius* | Stool | Central African Republic | 2 | NA | 0 |
| SAT508 | *Streptococcus salivarius* | Stool | Central African Republic | 3 | NA | 0 |
| SAT520 | *Streptococcus salivarius* | Stool | Central African Republic | 1 | NA | 0 |
| SAT526 | *Streptococcus salivarius* | Stool | Central African Republic | 3 | NA | 0 |
| SAT535 | *Streptococcus salivarius* | Stool | Central African Republic | 2 | NA | 0 |
| SAT536 | *Streptococcus salivarius* | Stool | Central African Republic | 1 | NA | 0 |
| SAT502 | *Streptococcus parasanguinis* | Stool | Central African Republic | 0 | NA | 0 |
| SAT504 | *Streptococcus parasanguinis* | Stool | Central African Republic | 1 | NA | 0 |

| | | | | | | |
|---|---|---|---|---|---|---|
| SAT506 | *Streptococcus parasanguinis* | Stool | Central African Republic | 0 | NA | 0 |
| SAT509 | *Streptococcus parasanguinis* | Stool | Central African Republic | 1 | NA | 0 |
| SAT511 | *Streptococcus parasanguinis* | Stool | Central African Republic | 0 | NA | 0 |
| SAT529 | *Streptococcus parasanguinis* | Saliva | Central African Republic | 0 | NA | 0 |
| SAT532 | *Streptococcus parasanguinis* | Saliva | Central African Republic | 1 | NA | 0 |
| SAT551 | *Streptococcus parasanguinis* | Saliva | Central African Republic | 0 | NA | 0 |
| SAT561 | *Streptococcus parasanguinis* | Saliva | Central African Republic | 0 | NA | 2 |
| SAT565 | *Streptococcus parasanguinis* | Saliva | Central African Republic | 0 | NA | 0 |
| SAT521 | *Streptococcus mitis* | Saliva | Central African Republic | 0 | NA | 2 |
| SAT522 | *Streptococcus mitis* | Saliva | Central African Republic | 0 | NA | 3 |
| SAT523 | *Streptococcus mitis* | Saliva | Central African Republic | 0 | NA | 2 |
| SAT524 | *Streptococcus mitis* | Saliva | Central African Republic | 0 | NA | 3 |
| SAT527 | *Streptococcus mitis* | Saliva | Central African Republic | 0 | NA | 3 |
| SAT534 | *Streptococcus mitis* | Saliva | Central African Republic | 0 | NA | 3 |
| SAT538 | *Streptococcus mitis* | Saliva | Central African Republic | 0 | NA | 3 |
| SAT563 | *Streptococcus mitis* | Saliva | Central African Republic | 0 | NA | 2 |
| SAT568 | *Streptococcus mitis* | Saliva | Central African Republic | 0 | NA | 1 |

**Supplementary Table 2.** Strain used to generate the spent medium ("donor" strains)

| **Strain ID** | **Strain ID2** | **Source** | **Bacterial species** | **Niche overlap** |
|---|---|---|---|---|
| This Study | AFB099 | Human feces, Central African Republic | *Coprococcus comes* | YES |
| This Study | AFB150 | Human feces, Central African Republic | *Blautia intestinalis* | YES |
| This Study | AFB159 | Human feces, Central African Republic | *Ruminococcus lactaris* | YES |
| This Study | AFB180 | Human feces, Central African Republic | *Blautia wexlerae* | YES |
| This Study | AFB204 | Human feces, Central African Republic | *Flavonifractor plautii* | YES |
| This Study | AFB210 | Human feces, Central African Republic | *Bacteroides ovatus* | YES |
| This Study | AFB226 | Human feces, Central African Republic | *Coprococcus eutactus* | YES |

| | | | | |
|---|---|---|---|---|
| This Study | AFB290 | Human feces, Central African Republic | *Blautia glucerasea* | YES |
| This Study | AFB426 | Human feces, Central African Republic | *Ruminococcus bromii* | YES |
| This Study | AFB475 | Human feces, Central African Republic | *Blautia luti* | YES |
| This Study | AFB478 | Human feces, Central African Republic | *Blautia caecimuris* | YES |
| This Study | AFB481 | Human feces, Central African Republic | *Paraclostridium bifermentans* | YES |
| This Study | AFB493 | Human feces, Central African Republic | *Blautia obeum* | YES |
| This Study | AFB594 | Human feces, Central African Republic | *Odoribacter splanchnicus* | YES |
| This Study | EBT030 | Human feces, Switzerland | *Bacteroides intestinalis* | YES |
| This Study | EBT048 | Human feces, Switzerland | *Bacteroides faecis* | NO |
| This Study | EBT058 | Human feces, Switzerland | *Bacteroides caccae* | YES |
| This Study | EBT067 | Human feces, Switzerland | *Mediterraneibacter faecis* | YES |
| This Study | EBT072 | Human feces, Switzerland | *Anaerobutyricum hallii* | NO |

| | | | | |
|---|---|---|---|---|
| This Study | EBT151 | Human feces, Switzerland | *Anaerobutyricum hallii* | NO |
| This Study | EBT260 | Human feces, Switzerland | *Ruminococcus faecis* | YES |
| DSM3319 | PV008 | Human feces | *Anaerostipes hadrus* | YES |
| CCUG69734 | PV015 | Human blood, Sweden | *Anaerotruncus colihominis* | YES |
| ATCC29236 | PV025 | NA | *Blautia coccoides* | YES |
| DSM27629 | PV027 | Human feces, Republic of Korea | *Blautia faecis* | YES |
| DSM20583 | PV037 | Human feces | *Blautia hansenii* | YES |
| DSM10518 | PV067 | Rumen of suckling lamb, France | *Blautia schinkii* | YES |
| DSM105336 | PV086 | Wild boar, Canada | *Lacrimispora celerecrescens* | YES |
| ATCC25537 | PV096 | Calf rumen | *Enterocloster clostridioformis* | YES |
| DSM108250 | PV111 | Human feces, Austria | *Clostridium symbiosum* | YES |
| ATCC27761 | PV119 | NA | *Coprococcus catus* | YES |
| DSM15684 | PV158 | Human feces, USA | *Eubacterium ramulus* | YES |
| DSM108070 | PV200 | Human feces, Scotland | *Roseburia inulinivorans* | NO |

**Supplementary Tables 3.** Mice related measurement and statistics

**CFU/mL in feces of mice treated with lysin or PBS control**

| Day | Control (mean ± SD) | Lysin (mean ± SD) | p-value |
|---|---|---|---|
| 21 | 1745.4 ± 1119.8 | 1472.2 ± 1350.5 | 0.589 |
| 22 | 2031.2 ± 1699.2 | 1500.3 ± 1268.3 | 0.485 |
| 23 | 1640.5 ± 1431.6 | 455.6 ± 214.6 | 0.093 |
| 24 | 1466.8 ± 1121.4 | 603.1 ± 218.9 | 0.132 |

*Values are mean ± SD. p-values were calculated using Mann–Whitney U test. (N=6).

**CFU/mL in small-intestinal segments of mice treated with lysin or PBS control**

| Organ | Control (mean ± SD) | Lysin (mean ± SD) | p-value |
|---|---|---|---|
| Duodenum | 207.9 ± 62.5 | 203.0 ± 153.9 | 0.093 |
| Ileum | 795.4 ± 834.3 | 237.4 ± 90.8 | 0.132 |
| Jejunum | 674.7 ± 773.1 | 244.0 ± 181.4 | 0.065 |

*Values are mean ± SD. p-values were calculated using Mann–Whitney U test. (N=6).

**Trend analyses of fecal CFU counts**

| Group | Slope (CFU/day) | SE | p-value |
|---|---|---|---|
| Control | -122.7 | 239.3 | 0.613 |
| Lysin | -365.2 | 170.5 | 0.044 |

* Linear regression was applied to replicate-level data across days (D21–D24). The slope indicates the estimated daily change in CFU/ml. A negative slope indicates a downward trend. Significance determined by regression p-value.

**Raw CFU/mL in feces**

| Day | Group | Rep1 | Rep2 | Rep3 | Rep4 | Rep5 | Rep6 |
|---|---|---|---|---|---|---|---|
| 21 | Control | 402.7 | 937.2 | 2040.2 | 2125.6 | 3599.9 | 1366.9 |
| 21 | Lysin | 1027.7 | 1574.1 | 859.4 | 4091.6 | 259.2 | 1021.2 |
| 22 | Control | 1131.3 | 334.9 | 5154.7 | 2544.1 | 1818.7 | 1203.8 |
| 22 | Lysin | 653.3 | 965.1 | 756.9 | 3915.6 | 1907.9 | 802.8 |
| 23 | Control | 615.3 | 167.3 | 1963.0 | 1065.2 | 4200.6 | 1831.6 |
| 23 | Lysin | 258.2 | 436.8 | 634.0 | 713.4 | 160.7 | 530.5 |
| 24 | Control | 571.4 | 514.3 | 908.6 | 1039.7 | 3073.2 | 2693.4 |
| 24 | Lysin | 602.2 | 482.5 | 889.0 | 742.8 | 254.0 | 647.8 |

**Raw CFU/mL in intestinal organs**

| Organ | Group | Rep1 | Rep2 | Rep3 | Rep4 | Rep5 | Rep6 |
|---|---|---|---|---|---|---|---|
| Duodenum | Control | 295.9 | 127.3 | 197.6 | 177.4 | 267.5 | 181.5 |
| Duodenum | Lysin | 75.8 | 154.4 | 55.6 | 287.2 | 176.6 | 468.3 |

| | | | | | | | |
|---|---|---|---|---|---|---|---|
| Ileum | Control | 41.1 | 258.3 | 2399.1 | 796.3 | 608.9 | 669.0 |
| Ileum | Lysin | 242.4 | 194.2 | 231.3 | 297.5 | 96.1 | 362.7 |
| Jejunum | Control | 230.3 | 422.8 | 270.0 | 236.6 | 673.8 | 2214.7 |
| Jejunum | Lysin | 597.0 | 176.6 | 241.9 | 211.9 | 155.8 | 80.8 |

**Mouse body weight (g) at different days post-arrival**

| MouseID | Treatment | Day0 | Day7 | Day14 | Day21 | Day22 | Day23 | Day24 |
|---|---|---|---|---|---|---|---|---|
| XB_0119 | + Lysin | 10.2 | 11.6 | 12.6 | 14.0 | 14.6 | 14.7 | 14.7 |
| XB_0120 | + Lysin | 10.4 | 11.9 | 13.4 | 15.2 | 15.5 | 15.9 | 16.0 |
| XB_0121 | PBS control | 10.4 | 12.0 | 13.0 | 14.3 | 14.8 | 14.9 | 14.8 |
| XB_0122 | PBS control | 9.6 | 11.2 | 12.2 | 14.3 | 13.7 | 13.5 | 13.8 |
| XB_0123 | PBS control | 10.2 | 10.8 | 12.2 | 14.0 | 14.0 | 14.0 | 14.3 |
| XB_0124 | PBS control | 9.9 | 11.7 | 12.3 | 13.2 | 14.1 | 14.4 | 14.3 |
| XB_0125 | + Lysin | 9.6 | 10.9 | 12.5 | 14.0 | 14.3 | 14.5 | 14.0 |
| XB_0126 | + Lysin | 9.7 | 12.1 | 13.2 | 14.9 | 15.2 | 15.0 | 14.9 |
| XB_0127 | + Lysin | 9.7 | 11.9 | 13.1 | 14.6 | 15.0 | 14.9 | 15.1 |
| XB_0128 | + Lysin | 10.4 | 11.7 | 13.6 | 14.6 | 15.1 | 15.5 | 15.6 |
| XB_0129 | PBS control | 9.9 | 12.0 | 13.9 | 15.9 | 16.1 | 16.6 | 17.1 |

| | | | | | | | | |
|---|---|---|---|---|---|---|---|---|
| XB_0130 | PBS control | 9.6 | 11.7 | 13.8 | 15.8 | 16.3 | 16.6 | 16.9 |
| Mean ± SD (Control vs Lysin) | | 9.93 ± 0.32 vs 10.00 ± 0.37 (p=0.744) | 11.57 ± 0.48 vs 11.68 ± 0.42 (p=0.808) | 12.90 ± 0.79 vs 13.07 ± 0.44 (p=0.575) | 14.58 ± 1.06 vs 14.55 ± 0.48 (p=0.936) | 14.83 ± 1.12 vs 14.95 ± 0.43 (p=0.589) | 15.00 ± 1.32 vs 15.08 ± 0.52 (p=0.630) | 15.20 ± 1.43 vs 15.05 ± 0.70 (p=0.810) |

Note: Values are presented as individual mouse measurements and group means ± SD. p-values are from Mann–Whitney U tests (two-sided).

**Mouse body weight gain vs Day 0 (g)**

| MouseID | Treatment | Gain at D21 | Gain at D22 | Gain at D23 | Gain at D24 |
|---|---|---|---|---|---|
| XB_0119 | + Lysin | 3.8 | 4.4 | 4.5 | 4.5 |
| XB_0120 | + Lysin | 4.8 | 5.1 | 5.5 | 5.6 |
| XB_0121 | PBS control | 3.9 | 4.4 | 4.5 | 4.4 |
| XB_0122 | PBS control | 4.7 | 4.1 | 3.9 | 4.2 |
| XB_0123 | PBS control | 3.8 | 3.8 | 3.8 | 4.1 |
| XB_0124 | PBS control | 3.3 | 4.2 | 4.5 | 4.4 |
| XB_0125 | + Lysin | 4.4 | 4.7 | 4.9 | 4.4 |
| XB_0126 | + Lysin | 5.2 | 5.5 | 5.3 | 5.2 |

| | | | | | |
|---|---|---|---|---|---|
| XB_0127 | + Lysin | 4.9 | 5.3 | 5.2 | 5.4 |
| XB_0128 | + Lysin | 4.2 | 4.7 | 5.1 | 5.2 |
| XB_0129 | PBS control | 6.0 | 6.2 | 6.7 | 7.2 |
| XB_0130 | PBS control | 6.2 | 6.7 | 7.0 | 7.3 |
| Mean ± SD (Control vs Lysin) | | 4.65 ± 1.21 vs 4.55 ± 0.51 (p=0.873) | 4.90 ± 1.23 vs 4.95 ± 0.42 (p=0.422) | 5.07 ± 1.42 vs 5.08 ± 0.35 (p=0.468) | 5.27 ± 1.54 vs 5.05 ± 0.49 (p=0.467) |

Note: Values are presented as individual mouse measurements and group means ± SD. p-values are from Mann–Whitney U tests (two-sided).
